# In Vivo Mutagenesis of a Ketosynthase Domain Uncovers Productivity and Specificity Control in Modular Polyketide Synthases

**DOI:** 10.1101/2025.09.26.678738

**Authors:** Jingyi Hu, Susanna Kushnir, Marius Brandenburger, Frank Schulz

## Abstract

Ketosynthase domains govern chain transfer and substrate selectivity in modular polyketide synthases (PKS), yet their functional tunability in native contexts remains poorly understood. We performed phylogenetically guided mutagenesis of the KS5 domain from the *Streptomyces cinnamonensis* monensin PKS and evaluated 72 variants *in vivo* across wild-type and reductive-loop-null backgrounds. This revealed discrete active-site motifs that control productivity, redox-state specificity, and extender-unit selection, functions traditionally ascribed to other PKS domains. AlphaFold3 structural mapping linked these motifs to substrate-tunnel and catalytic-core features, providing a mechanistic basis for the observed phenotypes. Our findings demonstrate that KS domains can be rationally re-tuned to overcome productivity bottlenecks and alter specificity in intact PKSs, offering a route to improved yields and expanded chemical diversity in engineered polyketides.

## Introduction

Polyketides represent one of the most structurally diverse classes of natural products, producing clinically important compounds such as erythromycin, rapamycin, and numerous anticancer agents ^1,2^. These molecules are assembled by type I polyketide synthases (PKS), complex modular megaenzymes that operate as molecular assembly lines. The modular architecture of these systems, where each module typically contains ketosynthase (KS), acyltransferase (AT), and acyl-carrier protein (ACP) domains, creates the structural foundation for polyketide diversity. Optional reductive domains, including ketoreductase (KR), dehydratase (DH), and enoylreductase (ER), further modify nascent β-keto groups, generating the varied oxidation patterns that distinguish different polyketide scaffolds ^3^.

Despite decades of research into PKS engineering, efforts to create novel polyketides through domain modifications consistently encounter a critical bottleneck: the ketosynthase domains act as stringent gatekeepers that severely restrict the passage of non-natural intermediates through the assembly line ^4–8^. This gatekeeper function becomes particularly problematic when upstream domains are engineered to produce modified substrates, as the downstream KS domains often reject these altered intermediates, leading to dramatically reduced product titers or complete pathway shutdown ^9^.

The monensin biosynthetic system in *Streptomyces cinnamonensis* provides an ideal platform for investigating KS domain plasticity and specificity. Monensin is assembled by a cis-AT PKS consisting of a loading module and 12 elongation modules, followed by extensive post-PKS tailoring ^10^. Previous engineering efforts have revealed that modifications to reductive domains consistently create bottlenecks at downstream KS domains ^9,11^. Recently, a single consensus-guided mutation in the KS5 domain of the monensin PKS yielded an increase by two orders of magnitude in polyketide titers, proving that targeted mutagenesis of KS active sites can unlock hidden catalytic potential ^12^.

Building on this, we hypothesized that systematic exploration of sequence variations could reveal the molecular determinants of KS specificity and activity. However, the experimental reality of PKS mutagenesis imposes severe constraints on this approach. Unlike smaller enzymes that can be subjected to high-throughput screening methods in a host like *E. coli*, the complexity and size of modular PKS systems limit researchers to constructing relatively small libraries of targeted variants ^11,13–15^. This experimental hurdle necessitates focused mutation design strategies to maximize the information gained from each constructed variant. In full-length PKS mutagenesis, randomization as used in directed enzyme evolution cannot be routinely employed ^16^.

To address this challenge, we first employed multiple sequence alignment (MSA) of evolutionarily related polyether-forming PKS systems to identify candidate motifs that correlate with specific catalytic traits. Next, we employed a platform for the medium-throughput mutagenesis of PKS inside *Streptomyces* spp., as recently described by us ^17^.

Our approach involved systematically introducing motifs identified through the MSA into the KS5 domain of the monensin PKS and testing their effects in three distinct recipients: the wild-type KS5 strain producing premonensin A and B, a KR4-null mutant producing keto-premonensin derivatives, and a DH4-null mutant producing hydroxy-premonensin derivatives. This experimental design allowed us to examine both the impact of specific mutations on redox substrate specificity and their effect on catalytic efficiency across distinct biosynthetic contexts.

## Results and Discussion

### Phylogenetic analysis and mutation design

Due to the size and complexity of modular PKSs, comprehensive mutational screening is impractical. Each variant demands labor-intensive cultivation and analysis, so our study of KS5 domain plasticity was limited to dozens of variants rather than the thousands typical of directed evolution ^18–20^. These constraints necessitated a strategic approach to mutation selection. Encouraged by our previous study ^12^, here we decided to continue using *Streptomyces. cinnamonsis A495* (*S. cinnamonsis* ATCC 15413 ΔmonCIΔmonBI/II) ^21^ as our model system.

First, to minimize background noise in the alignment, we selected five evolutionarily related polyether-forming PKSs: monensin, nanchangmycin, lasalocid, nigericin, and salinomycin. These systems share 61%–100% sequence identity (Matrix distance data available in ZENODO at 10.5281/zenodo.4478685, preview in Figure S1) enabling robust alignment and identification of variable regions as candidate motifs. An expansion of the set of PKS gene clusters in the alignment led to the commonly observed clustering of KS domains with their respective gene cluster^22^. KS domain sequences were then aligned using ClustalW ^23^, and a phylogenetic tree was generated with the Maximum Likelihood method (data available in ZENODO at 10.5281/zenodo.4478685, preview in Figure S2) ^24^.

The phylogenetic analysis revealed three sub-clades (Figure 1A), each predominantly associated with a distinct redox state of the incoming substrate: the H-clade (β-hydroxy, green), A-clade (β-aliphatic, blue), and O-clade (β-olefinic, purple). This is reminiscent of that observed in trans-AT PKSs, where KS domains cluster according to the redox state of their incoming intermediates, reflecting intrinsic redox substrate specificity ^27–32^.

**Figure 1:**
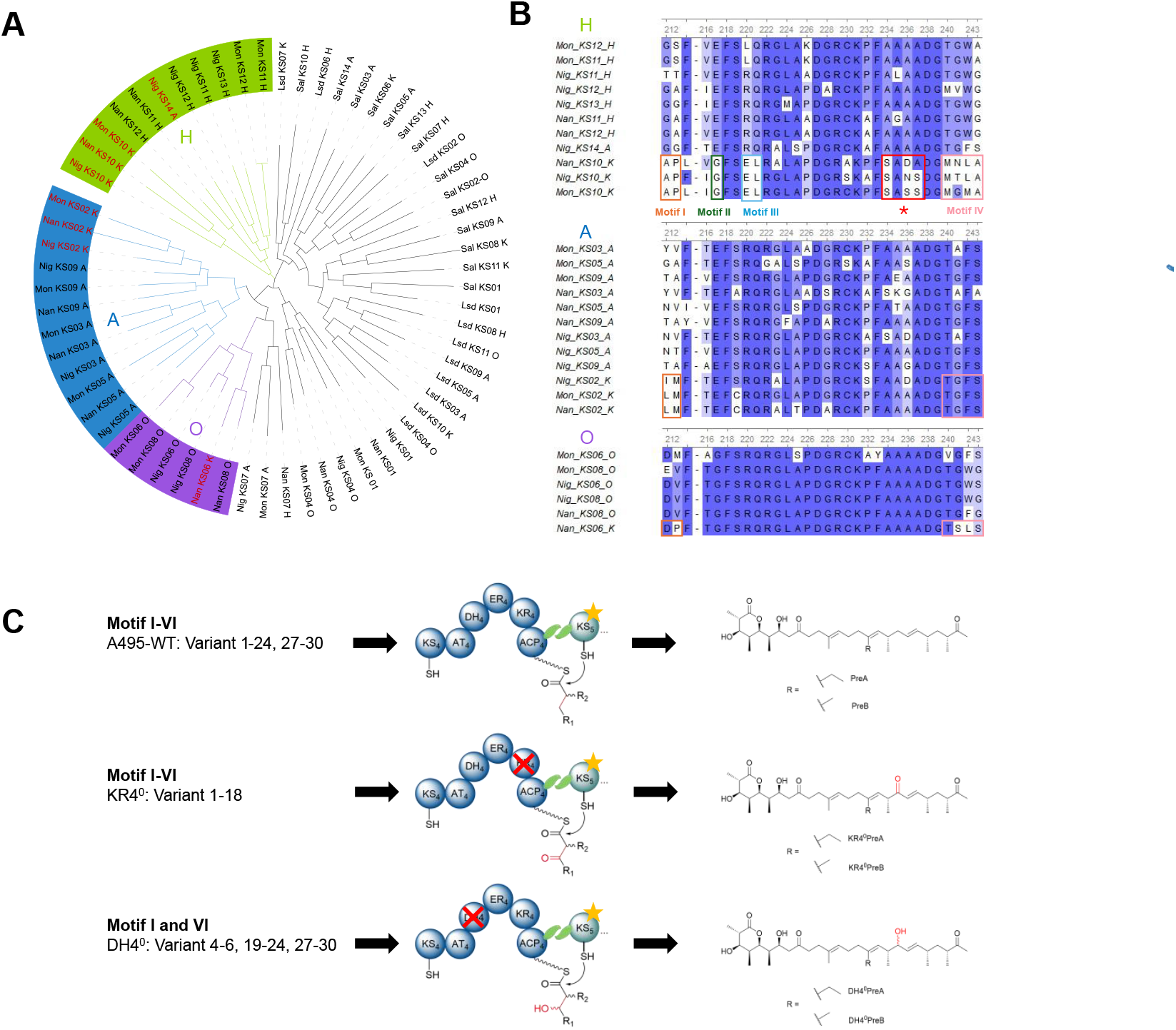
(A) Cladogram of KS domains from five closely related gene clusters (Monensin [Mon], Nanchangmycin [Nan], Lasalocid [Lsd], Nigericin [Nig], and Salinomycin [Sal]). The phylogenetic tree was inferred in MEGA7 using the Maximum Likelihood method with the JTT model of amino acid substitution, uniform rates among sites, partial deletion of gaps (30% cutoff), and branch support assessed with 1000 bootstrap replicates ^25^. KS numbering corresponds to the module number in which the KS domain resides, while the suffix indicates the redox state of the incoming substrate: _A for β-aliphatic, _K for β-keto, _H for β-hydroxy, _O for β-olefinic thioesters. For example, Mon_KS02_K denotes the KS domain in module 2 of the monensin PKS, which accepts a β-keto thioester. KS_01 designates the KS domain that transfers the acetyl starter unit and is therefore not labelled with a letter at the end. The three sub-clades that correlate with an apparent redox specificity of the corresponding KS domain were highlighted in green (H-hydroxy), blue (A-aliphatic), and purple (O-Olefinic), outliers of each sub-clade were highlighted in red. (B) MSA within each subclade identified four motifs (Motif I-IV) that corresponded with redox substrate specificity, plus one previously identified productivity-enhancing motif marked by an asterisk (full sequence alignments see Figure S3, data available in ZENODO at 10.5281/zenodo.4478685). Numbers above the sequence represent the column number of the alignment. Noteworthy is that all identified motifs are located near to each other, revealing an apparent connection between the sites. (C) Mutations in motifs I–IV were introduced into the KS5 domain of three recipients: A495-WT, KR4^0^ and DH4^0^. Due to the promiscuity of the AT5 domain ^26^, both methylmalonyl-CoA and ethylmalonyl-CoA can be incorporated in module 5, yielding products A and B. For the A495-WT recipient strain, producing premonensin A (preA) and B (preB), 28 variants were constructed. For the KR4-null (KR4^0^) recipient strain, 18 variants were constructed. In this recipient, the KR4 domain is inactivated, leading to the production of keto-premonensin derivatives KR4^0^PreA and KR4^0^PreB. For the DH4-null (DH4^0^) recipient strain, 13 variants were constructed. Here, the DH domain in module 4 is inactivated, resulting in the hydroxy-premonensin derivatives DH4^0^PreA and DH4^0^PreB.

Within each clade, we also identified several “outlier” KS domains that deviated from the typical substrate preference of their phylogenetic neighbors. Retrospective analysis indicated that these outliers originally processed intermediates with alternative redox states but adapted to β-keto substrates following inactivation of an upstream reductive module. This observation provided the key insight for our mutagenesis strategy: by comparing outlier sequences with the clade consensus, we can identify specific motifs potentially conferring altered substrate specificity.

This analysis yielded four candidate motifs containing residues that deviated from clade consensus patterns (Motifs I–I; Figure 1B), as well as a previously identified productivity-enhancing motif (Figure 1B, red box with asterisk) ^12^. We were inspired by the concept of saturation mutagenesis with a restricted amino acid set and systematically assessed the effects of alternative residues at individual sites, both singly and in selected combinations^33^. We systematically tested these variations in three recipient strains: *S. cinnamonensis* A495 recipient strain (A495-WT) producing premonensin A and B (PreA and PreB), a KR4-null mutant (KR4^0^) producing keto-premonensin derivatives (KR4^0^PreA and KR4^0^PreB), and a DH4-null mutant (DH4^0^) producing hydroxy-premonensin derivatives (DH4^0^PreA and DH4^0^PreB). To control background mutagenesis effects, we reconstituted wild-type KS5 domains in each recipient, generating: A495-KS5-wild type (A495-rWT), KR4^0^-KS5-reconstituted WT (KR4^0^-rWT), and DH4^0^-KS5-reconstituted WT (DH4^0^-rWT). This experimental design allowed us to assess whether specific mutations could redirect KS5 toward different substrate redox preferences while simultaneously monitoring effects on overall productivity.

### MSA guided approach enhances polyketide productivity

Twenty-eight sets of mutations were introduced into the A495-WT recipient strain, and four clones of each variant were fermented and analyzed by high-performance liquid chromatography–mass spectrometry (HPLC–MS) (Figure 2). The observed clone-to-clone variability is a typical trait of *Streptoymces* spp., causing substantial experimental error ^34^. Therefore, we focused our discussion on results that showed clear differences and with small error bars. Despite the relatively small number of clones per variant, outliers were removed when clearly discernible.

**Figure 2:**
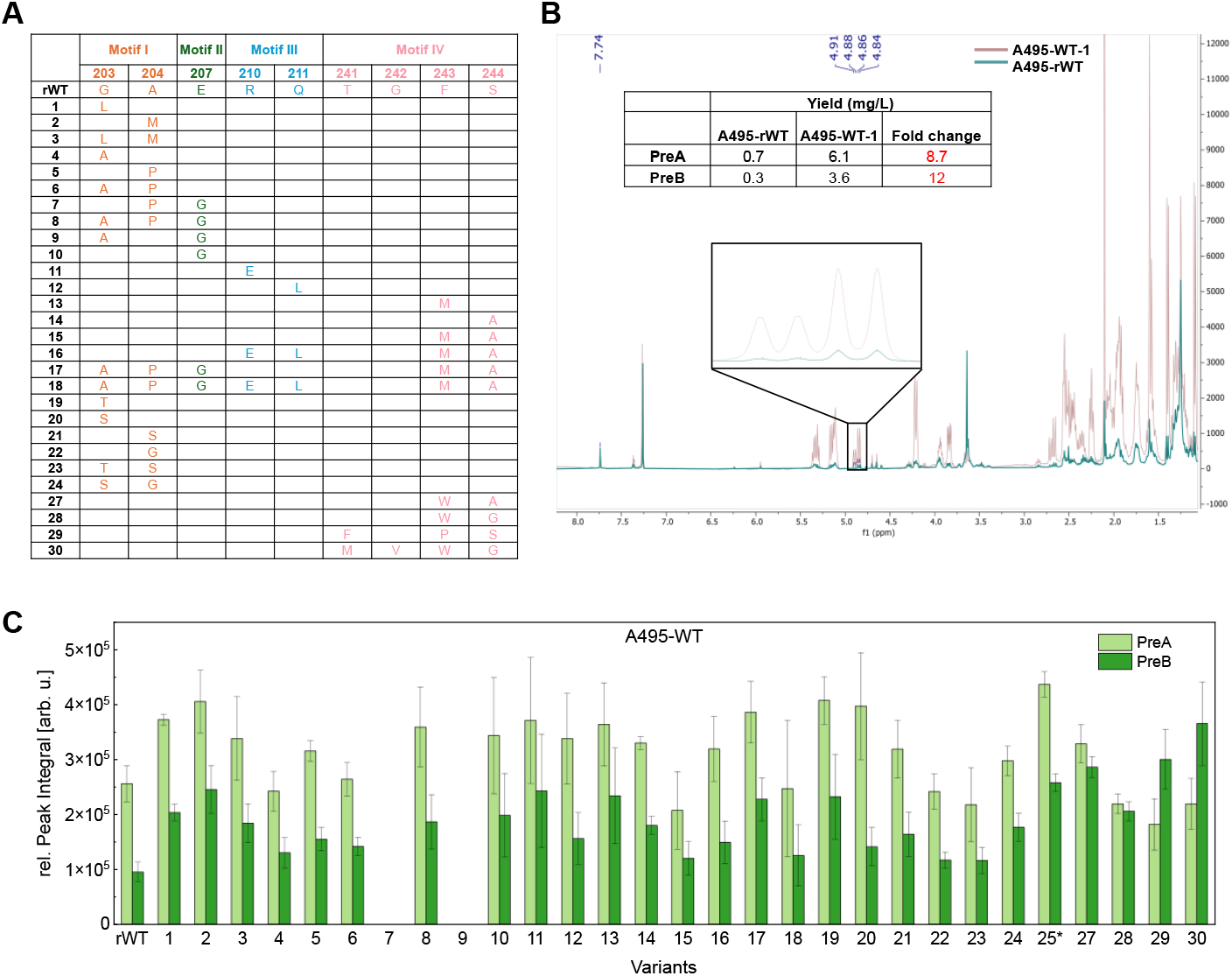
(A) 28 sets of motif I–IV mutations were introduced into the A495-WT recipient strain, with A495-rWT serving as control. The introduced template residues are shown in Figure S4. (B) Superposition of 1H-NMR spectra of clone 833 (A495-rWT, green) and clone 284 (A495-WT-1, brown). Absolute quantification of PreA and PreB was based on the H17 signals (PreB: 4.90 ppm, doublet; PreA: 4.85ppm, doublet), with 1,2,4,5-tetrachloronitrobenzene as an internal standard (7.74 ppm, singlet). Compared with clone 883 from A495-rWT, clone 284 from A495-WT-1 showed an 8.7-fold increase in PreA and a 12-fold increase in PreB. (C) Combined HPLC–MS peak integrals of adducts [M+Na]^+^, [M+H]^+^, [M+K]^+^, and [M+NH_4_]^+^ of PreA and PreB in A495-WT variants. Variant 25* was identified in our previous study, where it exhibited up to a 27-fold increase in productivity ^12^. All values were normalized to cell wet weight. Four independent clones of each variant were fermented, extracted, and analyzed using HPLC-MS. Error bars indicate standard deviation across clones.

Overall, the HPLC-MS-based screening showed large productivity variations across variants in A495-WT (Figure 2C). ariants 7 and 9 (Figure 2A and 2C) lost productivity entirely, whereas variant 8 maintained or even slightly improved its productivity. These results demonstrated the non-independence of individual mutations, with several showing strong antagonistic effects ^20^. The E207G substitution (motif II) or G203A/A204P (motif I) alone were tolerated, but combining E207G with either G203A or A204P abolished activity. Introducing all three substitutions together (E207G + G203A + A204P) rescued productivity, indicating a tight functional coupling between motifs I and II.

Four variants (A495-WT-1, -2, -17, -19, and -20) showed clear productivity enhancement. Compared with A495-rWT, titers of PreA increased ∼1.5-fold, while PreB increased 1.5–2.6-fold. As HPLC–MS has intrinsic limitations for absolute quantification, we validated these findings using quantitative NMR (qNMR) ^35^. In 1.5 L fermentations, qNMR revealed that a clone (A495-1) produced 8.7-fold more PreA and 12-fold more PreB than the rWT (Figure 2B), showing that HPLC–MS had underestimated the productivity gains, consistent with our previous work on premonensin ^12^.

Rather than altering substrate specificity as suggested by the MSA, certain mutations, such as G203L in variant A495-WT-1, enhanced catalytic efficiency of the KS5 domain. Such gain-of-productivity effects demonstrated that straightforward targeted mutations can improve yields in engineered PKS systems, providing a faster alternative to complex, long-term strain optimization.

### Motif III–IV mutations shift KS substrate specificity

To test whether mutations intended to enhance acceptance of β-keto substrates would alter KS5 specificity, mutation sets 1–18 (Figure 2A) were introduced into the KR4^0^-recipient strain. The resulting variants KR4^0^-1 to KR4^0^-18 were tested as described for the A495-WT variants (*vide supra*).

The results revealed a complex pattern of context-dependent effects (Figure 3). KR4^0^-1 and KR4^0^-2 both showed ∼2-fold increase in KR4^0^PreA/B production, similar to the effects observed in A495-WT-1 and -2 with the same mutations. This indicated enhanced overall productivity rather than altered specificity. Similarly, KR4^0^-7 and KR4^0^-9 lost productivity entirely, mirroring the A495-WT variants and again pointing to global activity loss rather than substrate selectivity shift.

**Figure 3:**
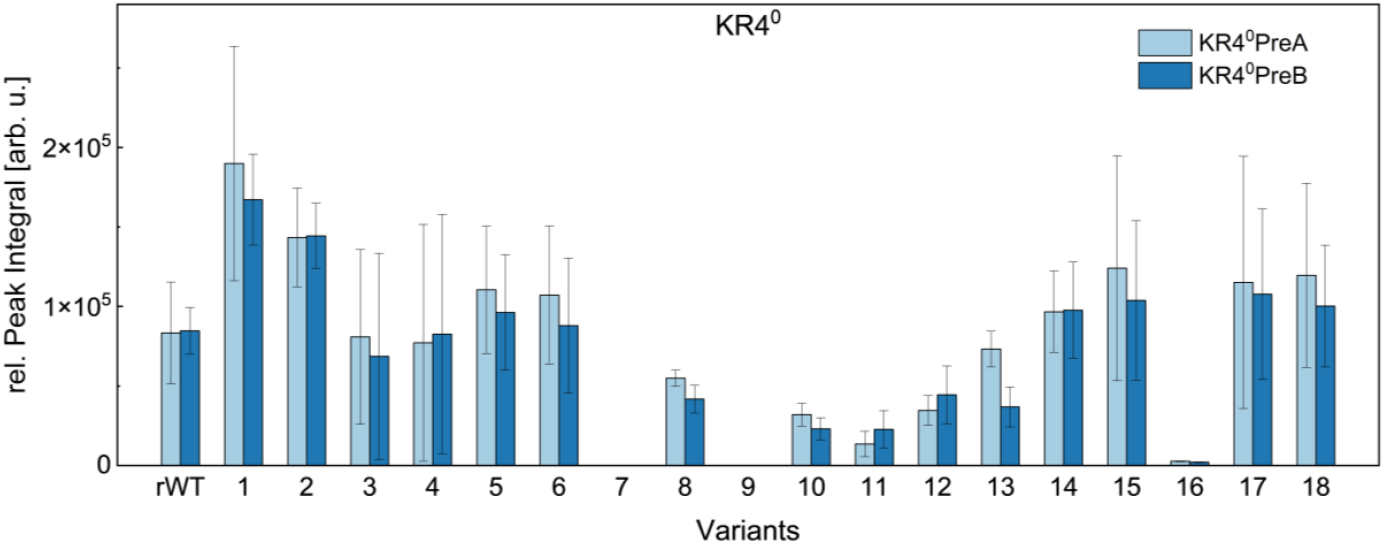
Combined HPLC–MS peak integrals of adducts [M+Na]^+^, [M+H]^+^, [M+K]^+^, and [M+NH_4_]^+^ of KR4^0^PreA and KR4^0^PreB in KR4^0^ variants. All values were normalized to cell wet weight. Four independent clones of each variant were fermented, extracted, and analyzed using HPLC-MS. Error bars indicate standard deviation across clones.

The strongest specificity effects emerged from mutations in motifs III and IV. Charge-reversing (R221E, KR4^0^-11 variant) and polarity-reversing (222L, KR4^0^-12 variant) substitutions in motif III markedly reduced keto-premonensin production while leaving premonensin biosynthesis unaffected. This pattern suggested that these mutations conferred stricter substrate specificity. The combination of motif III and IV mutations produced even stronger specificity shifts. The motif III & IV double mutant (KR4^0^-16 variant) almost abolished keto-premonensin production while maintaining premonensin levels, indicating that the engineered KS5 domain had developed a preference for fully reduced substrates over β-keto intermediates. The combined mutations across all four motifs (variant KR4^0^-18) fully restored keto-premonensin production, showing cooperative effects across all studied motifs, while the mutation of individual motifs exerts destructive effects.

These findings showed that the MSA-guided approach had indeed identified residues involved in substrate specificity determination, although their effects remain difficult to predict. In smaller enzymes, full saturation mutagenesis at the identified positions would be the next logical step, but the number of relevant sites and the properties of the *Streptomyces* host preclude this approach at present.

### Motif IV mutations inverts extender unit selectivity

The DH4^0^ variants provided a complementary test system, where dehydratase inactivation in module 4 leads to production of hydroxy-premonensin derivatives. This allowed us to examine whether MSA-identified mutations could impact acceptance of β-hydroxy substrates and thereby hydroxy-premonensin production, analogous to the effects found in the KR4^0^ system.

Mutation sets 4–6, 19–24, and 27–30 (Figure 2A) were introduced into the DH4^0^ variant. These mutations had different effects compared with the A495-WT recipient. For example, the G203T substitution (mutation 19) slightly enhanced premonensin production in A495-WT but had no significant effect on hydroxy-premonensin production in DH4^0^, whereas G203S (mutation 20) reduced productivity.

The most interesting observation in the DH4^0^ system came from mutations in motif IV (DH4^0^-27 to DH4^0^-30 variant), which inverted the extender unit selectivity. In the A495-rWT strain, the PreA/PreB ratio (ethylmalonyl-CoA vs. methylmalonyl-CoA incorporation) is typically 2.7. Motif IV mutations reduced this ratio to ∼1 or even inverted it, with A495-WT-30 producing predominantly PreB. This effect is even more significant in DH4^0^ variants, with variant DH4^0^-30 increasing DH4^0^PreB titers by fivefold (Figure 4A and 4B). These findings showed that KS domain mutations can influence extender unit selection, an activity traditionally attributed to AT domains. This reveals a previously underappreciated level of functional crosstalk between KS and AT domains within individual PKS modules ^36–38^.

**Figure 4:**
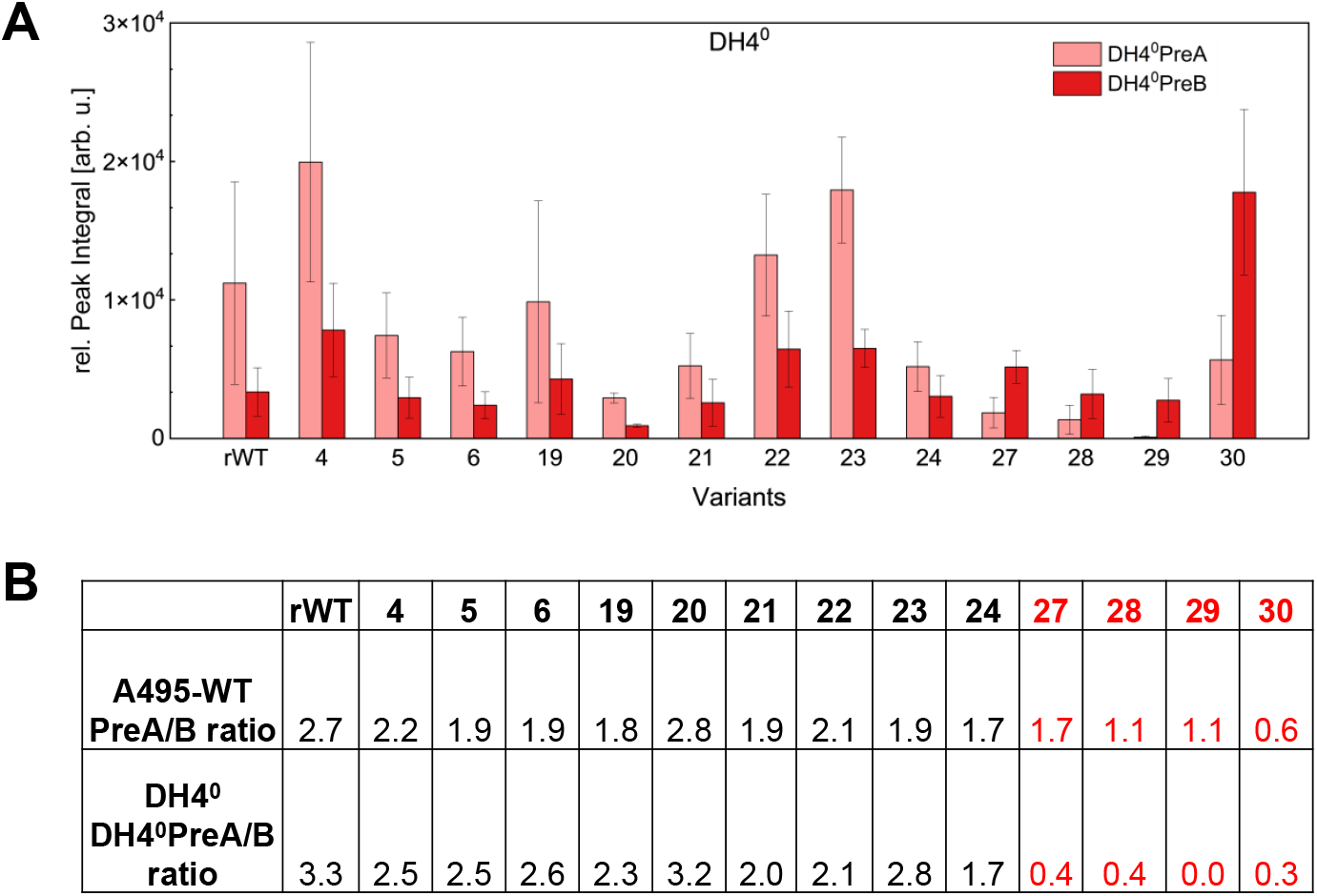
(A) Combined HPLC–MS peak integrals of adducts [M+Na]^+^, [M+K]^+^, and [M+NH_4_]^+^ of DH4^0^PreA and DH4^0^PreB in DH4^0^ variants. All values were normalized to cell wet weight. Four independent clones of each variant were fermented, extracted, and analyzed using HPLC-MS. Error bars indicate standard deviation across clones. (B) KS5 mutations 27–30 inverted extender unit selectivity in module 5 of A495-WT and DH40 variants. The PreA/PreB ratio in A495-WT decreased from 2.7 to 0.6–1.7, and the DH4^0^PreA/DH4^0^PreB ratio from 3.3 to 0.0– 0.3.

### Mechanistic Insights from Structural Analysis

To interrogate the molecular basis for the observed mutation effects, we generated AlphaFold3 models of the wild-type KS5 domain and mapped it to a FabF synthase bound to decanoyl-ACP (PDB: 8z5f) ^39,40^. FabF was chosen as reference because fatty acid synthase KSs and polyketide synthase KSs share a common evolutionary origin, with catalytic Cys-His-His triad and similar overall fold ^41–43^. This renders FabF suitable for the illustration and interpretation of the sequence-function-experiments described here, whereas the MonKS5/ACP5 dimer structure predicted by AlphaFold3 had confidence scores too low for reliable use.

The structural analysis revealed that the phylogenetically identified motifs map to distinct functional regions of the KS5 substrate binding tunnel. Motifs I and II residues (203, A204, and E207) cluster around the substrate binding cavity, positioned near the terminus of the nascent polyketide chain (Figure 5A and 5C). This localization explains why mutations in these regions primarily affect overall productivity rather than β-redox specificity, as they likely influence substrate orientation and binding during the catalytic cycle ^39^.

**Figure 5:**
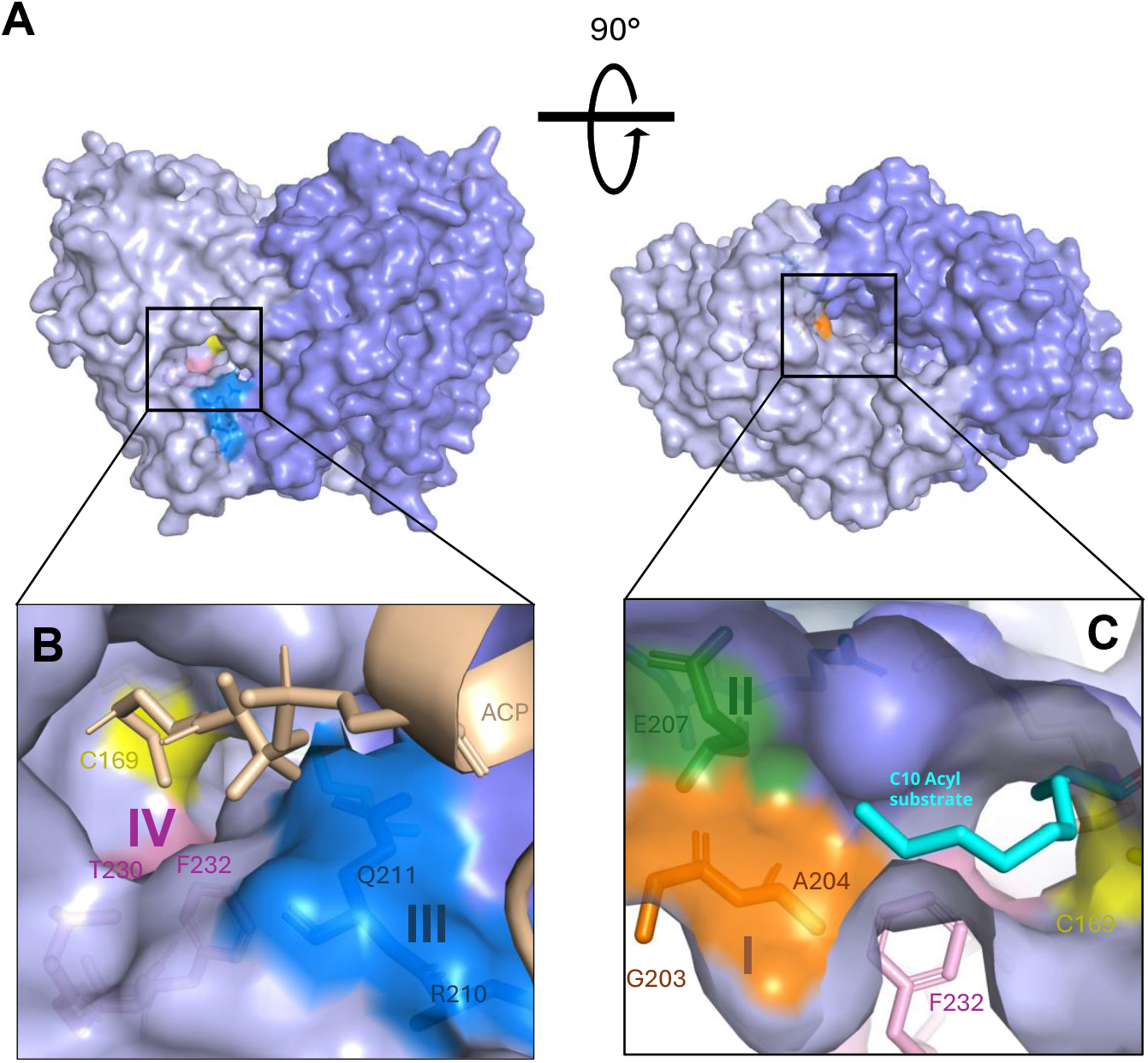
(A) AlphaFold3-predicted wild-type MonKS5 homodimer (monomers shown in light purple and slate). (B) Predicted KS5 WT model aligned with the FabF–decanoyl-ACP complex (PDB ID: 8Z5F, wheat). The FabF structure is not shown for clarity. The ACP phosphopantetheinyl (pPant) arm is shown as wheat sticks. Mutated residues T230 and F232 (motif IV, light pink) form the floor of the substrate tunnel directly beneath the catalytic cysteine (Cys169, yellow). Residues R210 and Q211 (motif III, blue) are located at the side entrance of KS5, near the ACP domain. (C) Residues G203 and A204 (motif I, orange) together with E207 (motif II, green,) shape the substrate tunnel at the homodimer interface. Superimposition of the substrate mimic decanone (cyan) from the FabF complex onto the MonKS5 model places the C10 atom in close proximity to these residues. This leaves them distant to the β-carbon atom of the nascent polyketide chain, hence an impact on the redox state is not to be expected (in line with the experimental observation but not with the MSA result). Yet, these residues can interact with the terminus of the nascent polyketide and thus can influence substrate binding which apparently is aided by an increase in size and hydrophobicity for residues 203 and 204.

Motif III residues (R221 and Q222) are located at the site entrance of the binding tunnel, where they could influence inter-module interactions mediated by upstream and downstream ACPs (Figure 5A and 5B) ^44–48^.

Motif IV residues are positioned near the catalytic cysteine (C169) of KS5, forming part of the active site (Figure 5A and 5B). Mutations in this region are therefore expected to alter the shape and chemical environment of the active site during the catalysis, consistent with the observed effects on extender unit selection and redox pattern preferences.

### Cooperation of Motif IV and Motif V in Regulating Specificity and Productivity

Further inspection of the AlphaFold model of the KS5 domain led to residue M411 (motif V) as a putative gating residue. Located on a loop adjacent to motif IV and catalytic Cys169, it contributes to form another part of the active site (Figure 6B). Similar gating residues, with phenylalanine at the equivalent position, have been reported in FabF and DEBS KS6 ^49,50^. Phylogenetic analysis revealed sequence variability at this position: alanine in the O-clade (olefinic substrates), isoleucine in most other clades, phenylalanine in two ketoacyl acceptors, and methionine uniquely in MonKS5 (Figure S3).

**Figure 6:**
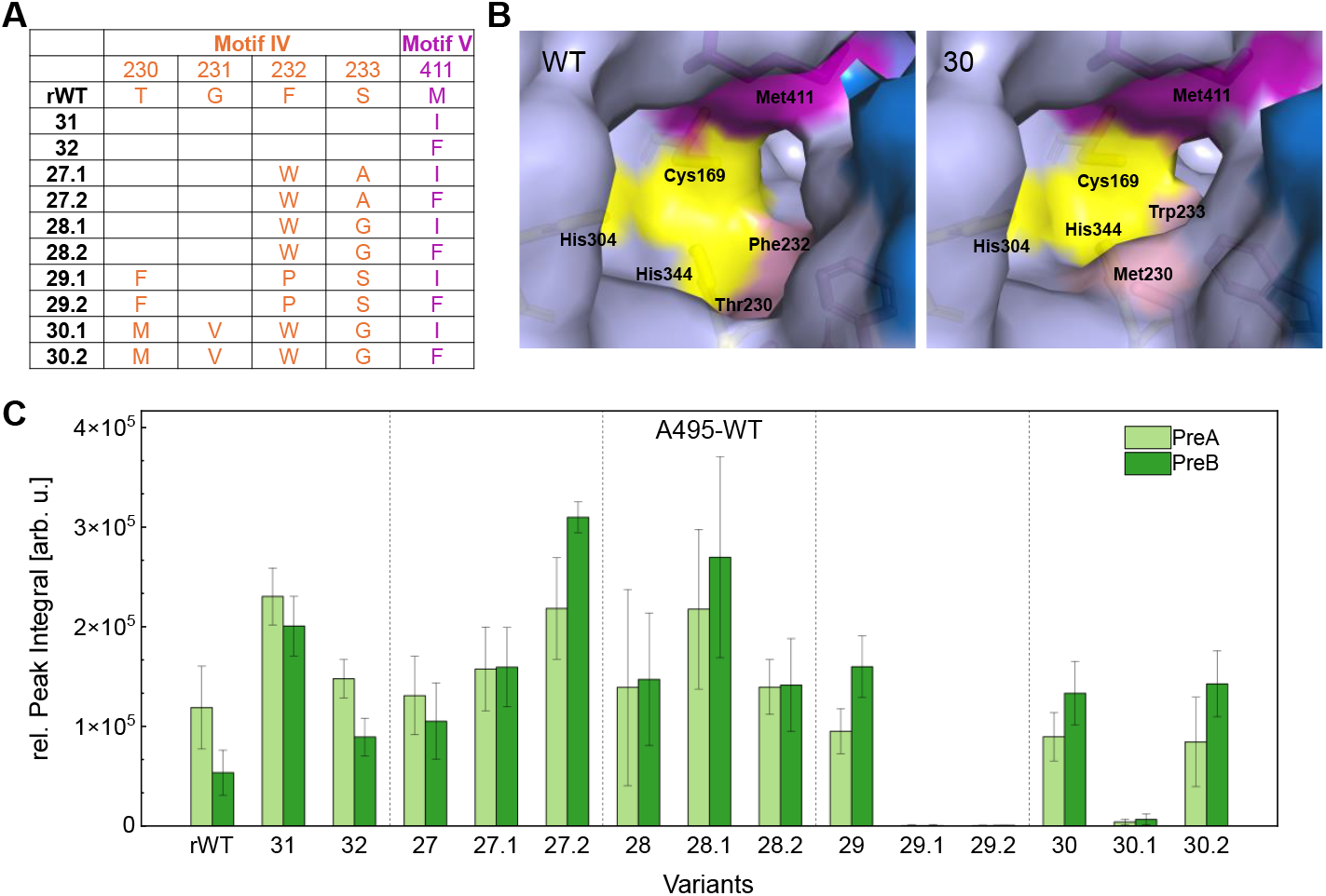
(A) Ten sets of motif IV–V mutations were introduced into A495-WT, with A495-rWT as the control. (B) AlphaFold3-predicted models of wild-type KS5 and KS5 with mutation set 30. Motif V (M411, purple) lies close to the Cys–His–His catalytic triad (yellow) and, together with motif IV (light pink), appears to form the core of the KS5 domain. It was therefore selected for further mutagenesis. (C)Combined HPLC–MS peak integrals of adducts [M+Na]^+^ and [M+K]^+^ of PreA and PreB in A495-WT variants. All values were normalized to cell wet weight. Four independent clones of each variant were fermented, extracted, and analyzed using HPLC-MS. Error bars indicate standard deviation across clones.

To probe its role, M411 was mutated alone and in combination with motif IV. Ten additional A495-WT variants were constructed and analyzed as described above (Figure 6A). Single motif V mutations enhanced productivity, with the consensus reversion M411I showing the strongest effect: variant 31 increased PreA and PreB titers 2- and 4-fold, respectively. ariant 27.2 displayed the best overall performance, nearly doubling PreA and increasing PreB sixfold, while variant 28.1 boosted PreA twofold and PreB fivefold. By contrast, Motif V mutations combined with mutation sets 29 or 30 were detrimental. All mutation 29-based variants almost lost KS activity entirely, while in mutation 30-based variants, M411I reduced production to trace levels, with only M411F maintaining productivity.

Structural modelling provided plausible explanations. In mutation 29-based variants, T230 was replaced with bulky hydrophobic phenylalanine while F232 was substituted with the smaller and rigid proline, changes likely to disrupt the spatial arrangement of the binding tunnel.

In mutation 30-based variants, all four motif IV residues were mutated, most to sterically more demanding residues. Here, T230M and S233W appear to constrict the active site, reducing the space available for substrate binding (Figure 7B). This is in line with the observed impact in DH4^0^30 variant, where the smaller extender unit methylmalonyl-CoA is favored over the wild-type preference for ethylmalonyl-CoA.

## Discussion

Our systematic, phylogenetically guided mutagenesis of the KS5 domain in the monensin PKS identified discrete motifs that control productivity, redox specificity, and extender-unit selection. These results support evolutionary models in which PKS diversity arises through module or domain duplication, followed by fine-tuning via point mutations or motif divergence ^22,51^. The most impactful substitutions either enhanced productivity or shifted extender-unit preference, demonstrating that natural PKS systems are not maximally optimized for catalytic efficiency and that gain-of-productivity mutations are achievable. These findings open the way to broadly applicable strategies for improving polyketide biosynthesis.

Despite these advances, several limitations remain. Mutation effects remain difficult to predict, even when guided by multiple sequence alignments, highlighting the need for high-resolution structural data and deeper mechanistic insight into PKS catalysis. In vivo characterization, though essential for capturing conformational flexibility and interdomain interactions, yields complex and context-dependent readouts compared with the cleaner outcomes of in vitro assays ^37,46,47,52–54^. Mutations that are beneficial in one genetic background may be neutral or detrimental in another, reflecting the sensitivity of KS domains to single-residue changes and the tight coupling of substrate binding, transacylation, and condensation. Furthermore, the integrated environment of the cell cannot yet be fully replicated in vitro, complicating the translation of engineering strategies to native contexts ^5,36,55–59^.

Even with these challenges, engineering modular PKSs remains both a formidable task and a promising opportunity. The same molecular complexity that makes these systems sensitive to sequence modifications also provides avenues for substantial performance gains. As mechanistic understanding deepens, the prospect of rationally designing PKS assembly lines with predictable properties and improved yields is increasingly within reach.

## Materials and methods

All protocols and oligonucleotides used are provided in the Supporting Information (SI).

### Bioinformatic analysis

KS domain sequences were retrieved from the Minimum Information about a Biosynthetic Gene cluster (MIBi) database, and phylogenetic trees were constructed using the Maximum Likelihood method (JTT model) with 1000 bootstrap replications in ME A7.0.26. Multiple sequence alignment was performed with ClustalW using the default settings in U ENE v48.1. Structural models were predicted with AlphaFold3 Beta and visualized in PyMOL. Complete phylogenetic tree, alignments, and model data are available in the SI (Figures S1–S3). The AlphaFold3-predicted model data are available at DOI: 10.5281/zenodo.4478685.

### KS mutagenesis

KS5 variants were generated as described by Bravo et al. ^12^. Briefly, mutants were constructed by overlap-extension PCR and cloned into a pKC1139-based vector using the GC-SLIC method ^17^. Constructs were integrated into the chromosome by conjugation-mediated homologous recombination, and mutations were verified by colony PCR. The oligonucleotides used are listed in Table S1.

### Fermentation, sample preparation, HPLC MS and qNMR analysis

Fermentation, sample preparation, and analytical procedures followed established methods^9,12^. Clones were grown on GYM agar, used to prepare TSB precultures (30 °C, 180 rpm). Preculture were further inoculated into 15 mL SM16 medium with 20g/L XAD16 resin. After 5 days, cells and resin were harvested, extracted with ethyl acetate, and dried extracts were dissolved in acetonitrile. HPLC–MS was performed on an Thermofisher Ultimate 3000 HPLC system with a NUCLEODUR® C18 column (Macherey&Nagel, 150/2, 1.8 μm) coupled to a Bruker Compact Q-TOF mass spectrometer (ESI, positive mode), and data were processed with MZmine 3 ^60^. For qNMR, 1.5 L cultures were extracted and purified by silica gel chromatography, and premonensin-containing fractions were reconstituted in 500 µL CDCl_3_ with 100 µL 1,2,4,5-tetrachloronitrobenzene (5mg/mL) as internal standard. Spectra were acquired on a Bruker A ANCE III HD 400 spectrometer (13C decoupling, 30 s relaxation delay) and processed in MestReNova 14. QNMR raw data are available at DOI: 10.5281/zenodo.4478685.

## Author Contributions

The authors thank Luca Marie Born for her contribution to the qNMR analyses of premonensin titers. M.B., J. H. and F.S. performed sequence alignment and designed the mutations. S.K. carried out mutagenesis, including oligonucleotide design and fermented bacteria. J.H. conducted experiments and data analysis, including testing the effects of mutagenesis on premonensin and derivative titers, AlphaFold analyses, and qNMR. The manuscript was written by J.H., and F.S.

## Acknowledgments

J. H. and F. S. acknowledge funding from the German Research Foundation (DF) through Research Training Group 2341 “Microbial Substrate Conversion (MiCon)”.

