## Supplementary Information for "In Vivo Mutagenesis of a Ketosynthase Domain Uncovers Productivity and Specificity Control in Modular Polyketide Synthases"

Raw and processed data for the bioinformatic analysis, the QNMR measurements and the AlphaFold3 models are available at the ZENODO repository under the following **DOI: 10.5281/zenodo.4478685**. The following sections contain preview representations for the bioinformatic analysis for the convenience of the reader.

### A) Bioinformatic analysis

| | Lsd_KS01 | Lsd_KS02_O | Lsd_KS03_A | Lsd_KS04_O | Lsd_KS05_A | Lsd_KS06_H | Lsd_KS07_K | Lsd_KS08_H | Lsd_KS09_A | Lsd_KS10_K | Lsd_KS11_O | Mon_KS02_K | Mon_KS03_A | Mon_KS04_O | Mon_KS05_A | Mon_KS06_O | Mon_KS07_A | Mon_KS08_O | Mon_KS09_A | Mon_KS10_K | Mon_KS11_H |
|---|---|---|---|---|---|---|---|---|---|---|---|---|---|---|---|---|---|---|---|---|---|
| Lsd_KS01 | 100% | 75% | 77% | 77% | 78% | 74% | 71% | 75% | 74% | 74% | 75% | 70% | 68% | 71% | 70% | 71% | 69% | 72% | 70% | 65% | 69% |
| Lsd_KS02_O | 75% | 100% | 75% | 79% | 76% | 76% | 72% | 77% | 78% | 73% | 79% | 73% | 70% | 72% | 71% | 71% | 69% | 73% | 72% | 65% | 72% |
| Lsd_KS03_A | 76% | 75% | 100% | 78% | 83% | 74% | 69% | 75% | 76% | 75% | 78% | 70% | 69% | 69% | 72% | 70% | 67% | 70% | 70% | 64% | 67% |
| Lsd_KS04_O | 77% | 79% | 78% | 100% | 80% | 75% | 72% | 77% | 78% | 76% | 81% | 72% | 73% | 74% | 73% | 73% | 69% | 75% | 74% | 67% | 71% |
| Lsd_KS05_A | 78% | 76% | 83% | 80% | 100% | 74% | 69% | 76% | 80% | 76% | 81% | 70% | 70% | 71% | 70% | 71% | 68% | 72% | 70% | 65% | 69% |
| Lsd_KS06_H | 74% | 76% | 74% | 75% | 74% | 100% | 74% | 72% | 73% | 72% | 75% | 69% | 68% | 72% | 70% | 69% | 66% | 69% | 73% | 65% | 71% |
| Lsd_KS07_K | 71% | 72% | 69% | 71% | 69% | 74% | 100% | 68% | 71% | 67% | 71% | 68% | 64% | 68% | 66% | 64% | 65% | 67% | 71% | 64% | 70% |
| Lsd_KS08_H | 74% | 77% | 75% | 77% | 76% | 72% | 68% | 100% | 77% | 73% | 80% | 71% | 68% | 69% | 68% | 70% | 66% | 72% | 69% | 64% | 69% |
| Lsd_KS09_A | 74% | 78% | 76% | 78% | 80% | 73% | 71% | 77% | 100% | 73% | 85% | 71% | 70% | 72% | 70% | 71% | 67% | 72% | 72% | 67% | 68% |
| Lsd_KS10_K | 73% | 72% | 75% | 75% | 76% | 72% | 67% | 73% | 73% | 100% | 75% | 69% | 66% | 68% | 66% | 67% | 66% | 69% | 67% | 65% | 66% |
| Lsd_KS11_O | 74% | 79% | 78% | 81% | 80% | 75% | 71% | 80% | 85% | 75% | 100% | 72% | 70% | 72% | 73% | 73% | 69% | 73% | 72% | 69% | 70% |
| Mon_KS02_K | 70% | 73% | 70% | 72% | 70% | 70% | 68% | 71% | 71% | 70% | 72% | 100% | 70% | 69% | 71% | 68% | 69% | 68% | 74% | 66% | 69% |
| Mon_KS03_A | 67% | 69% | 69% | 72% | 69% | 68% | 64% | 68% | 70% | 66% | 70% | 69% | 100% | 72% | 74% | 66% | 73% | 69% | 69% | 64% | 69% |
| Mon_KS04_O | 71% | 72% | 69% | 73% | 70% | 72% | 68% | 69% | 71% | 68% | 72% | 69% | 72% | 100% | 71% | 70% | 73% | 73% | 73% | 68% | 73% |
| Mon_KS05_A | 70% | 71% | 73% | 72% | 70% | 70% | 66% | 68% | 70% | 67% | 73% | 71% | 74% | 71% | 100% | 69% | 75% | 70% | 71% | 66% | 69% |
| Mon_KS06_O | 71% | 70% | 70% | 73% | 71% | 69% | 64% | 70% | 71% | 67% | 73% | 68% | 66% | 70% | 68% | 100% | 67% | 90% | 70% | 64% | 66% |
| Mon_KS07_A | 69% | 69% | 68% | 69% | 68% | 67% | 65% | 67% | 67% | 66% | 70% | 69% | 74% | 73% | 75% | 67% | 100% | 70% | 70% | 66% | 68% |
| Mon_KS08_O | 72% | 72% | 71% | 75% | 71% | 69% | 67% | 72% | 72% | 69% | 73% | 68% | 69% | 73% | 69% | 90% | 70% | 100% | 72% | 66% | 69% |
| Mon_KS09_A | 70% | 72% | 70% | 74% | 69% | 73% | 71% | 69% | 72% | 67% | 72% | 74% | 69% | 73% | 71% | 70% | 70% | 72% | 100% | 65% | 71% |
| Mon_KS10_K | 65% | 65% | 64% | 67% | 64% | 65% | 64% | 64% | 67% | 65% | 69% | 66% | 64% | 68% | 66% | 64% | 65% | 66% | 65% | 100% | 69% |
| Mon_KS11_H | 68% | 72% | 67% | 70% | 68% | 71% | 70% | 69% | 68% | 66% | 70% | 69% | 69% | 73% | 69% | 66% | 68% | 69% | 71% | 69% | 100% |
| Mon_KS12_H | 68% | 72% | 67% | 70% | 68% | 71% | 70% | 69% | 68% | 66% | 70% | 69% | 69% | 73% | 69% | 66% | 68% | 69% | 71% | 69% | 100% |
| Mon_KS_01 | 74% | 73% | 71% | 73% | 72% | 70% | 66% | 69% | 71% | 67% | 72% | 69% | 72% | 75% | 72% | 70% | 72% | 72% | 70% | 66% | 72% |
| Nan_KS01 | 72% | 69% | 71% | 72% | 72% | 71% | 66% | 70% | 72% | 66% | 75% | 68% | 70% | 72% | 71% | 70% | 68% | 71% | 70% | 65% | 71% |
| Nan_KS02_K | 72% | 68% | 70% | 72% | 70% | 68% | 68% | 69% | 71% | 68% | 70% | 75% | 68% | 70% | 69% | 67% | 68% | 67% | 73% | 64% | 67% |
| Nan_KS03_A | 70% | 70% | 69% | 71% | 70% | 69% | 65% | 68% | 70% | 66% | 70% | 68% | 81% | 72% | 71% | 69% | 71% | 71% | 70% | 65% | 71% |
| Nan_KS04_O | 73% | 73% | 71% | 73% | 73% | 72% | 67% | 71% | 72% | 68% | 73% | 70% | 70% | 81% | 69% | 68% | 67% | 70% | 70% | 64% | 71% |
| Nan_KS05_A | 70% | 72% | 72% | 72% | 71% | 71% | 69% | 69% | 71% | 67% | 71% | 69% | 73% | 73% | 77% | 70% | 71% | 71% | 73% | 66% | 69% |
| Nan_KS06_K | 68% | 69% | 70% | 72% | 69% | 68% | 65% | 68% | 71% | 66% | 72% | 68% | 71% | 72% | 70% | 73% | 69% | 76% | 70% | 65% | 69% |
| Nan_KS07_H | 68% | 69% | 68% | 71% | 66% | 71% | 65% | 68% | 67% | 67% | 69% | 68% | 70% | 67% | 70% | 66% | 72% | 67% | 71% | 61% | 67% |
| Nan_KS08_O | 69% | 70% | 72% | 73% | 72% | 68% | 65% | 68% | 72% | 67% | 72% | 69% | 71% | 71% | 70% | 73% | 69% | 76% | 74% | 64% | 68% |
| Nan_KS09_A | 70% | 73% | 69% | 72% | 70% | 72% | 69% | 70% | 71% | 68% | 71% | 75% | 70% | 75% | 73% | 72% | 70% | 73% | 79% | 65% | 71% |
| Nan_KS10_K | 66% | 67% | 68% | 69% | 66% | 67% | 64% | 66% | 66% | 66% | 66% | 65% | 66% | 67% | 66% | 64% | 63% | 66% | 66% | 72% | 68% |
| Nan_KS11_H | 72% | 74% | 74% | 74% | 73% | 74% | 69% | 71% | 73% | 70% | 75% | 70% | 69% | 71% | 70% | 69% | 68% | 71% | 70% | 64% | 74% |
| Nan_KS12_H | 73% | 74% | 74% | 74% | 73% | 75% | 71% | 74% | 73% | 71% | 76% | 70% | 69% | 71% | 71% | 70% | 69% | 72% | 71% | 66% | 77% |
| Nig_KS01 | 75% | 74% | 73% | 74% | 72% | 73% | 68% | 72% | 72% | 69% | 73% | 72% | 72% | 76% | 71% | 71% | 71% | 74% | 72% | 68% | 72% |
| Nig_KS02_K | 70% | 71% | 71% | 72% | 70% | 69% | 69% | 70% | 72% | 68% | 73% | 78% | 69% | 73% | 70% | 69% | 70% | 70% | 76% | 69% | 73% |
| Nig_KS03_A | 71% | 74% | 72% | 73% | 70% | 69% | 66% | 70% | 73% | 69% | 73% | 68% | 78% | 73% | 74% | 71% | 75% | 73% | 72% | 66% | 70% |
| Nig_KS04_O | 74% | 74% | 72% | 75% | 74% | 73% | 68% | 71% | 73% | 70% | 73% | 70% | 71% | 81% | 71% | 70% | 70% | 73% | 73% | 66% | 73% |
| Nig_KS05_A | 72% | 71% | 73% | 73% | 72% | 70% | 69% | 70% | 72% | 67% | 73% | 71% | 75% | 72% | 78% | 70% | 73% | 72% | 75% | 66% | 71% |
| Nig_KS06_O | 71% | 74% | 73% | 76% | 73% | 71% | 69% | 72% | 73% | 70% | 75% | 72% | 72% | 73% | 72% | 78% | 72% | 82% | 74% | 66% | 72% |
| Nig_KS07_A | 71% | 71% | 72% | 73% | 71% | 69% | 68% | 68% | 73% | 68% | 73% | 71% | 74% | 73% | 75% | 70% | 78% | 72% | 71% | 69% | 71% |
| Nig_KS08_O | 71% | 74% | 73% | 76% | 73% | 70% | 68% | 71% | 73% | 70% | 74% | 70% | 72% | 72% | 71% | 78% | 73% | 84% | 72% | 66% | 71% |
| Nig_KS09_A | 70% | 73% | 72% | 74% | 70% | 71% | 69% | 71% | 71% | 67% | 72% | 75% | 72% | 75% | 72% | 69% | 72% | 70% | 78% | 66% | 72% |
| Nig_KS10_K | 68% | 69% | 69% | 71% | 69% | 70% | 68% | 69% | 70% | 68% | 71% | 67% | 65% | 66% | 66% | 68% | 65% | 68% | 67% | 72% | 67% |
| Nig_KS11_H | 72% | 75% | 71% | 74% | 72% | 73% | 68% | 72% | 72% | 70% | 72% | 68% | 72% | 74% | 69% | 69% | 69% | 72% | 73% | 64% | 74% |
| Nig_KS12_H | 72% | 75% | 74% | 74% | 73% | 74% | 69% | 71% | 72% | 69% | 74% | 69% | 69% | 73% | 68% | 69% | 68% | 71% | 72% | 67% | 75% |
| Nig_KS13_H | 69% | 72% | 69% | 73% | 69% | 73% | 68% | 68% | 70% | 66% | 72% | 69% | 68% | 72% | 68% | 69% | 66% | 71% | 69% | 65% | 75% |
| Nig_KS14_A | 70% | 73% | 73% | 73% | 72% | 71% | 67% | 71% | 73% | 68% | 75% | 69% | 70% | 72% | 71% | 68% | 67% | 69% | 70% | 66% | 74% |
| Sal_KS01 | 76% | 71% | 69% | 68% | 70% | 67% | 68% | 67% | 72% | 67% | 72% | 68% | 68% | 71% | 71% | 65% | 68% | 66% | 67% | 62% | 67% |
| Sal_KS02-O | 71% | 79% | 71% | 74% | 73% | 69% | 69% | 72% | 73% | 69% | 74% | 69% | 69% | 70% | 68% | 67% | 67% | 70% | 69% | 64% | 72% |
| Sal_KS03_A | 71% | 75% | 75% | 72% | 75% | 73% | 69% | 72% | 73% | 68% | 75% | 67% | 68% | 68% | 68% | 67% | 67% | 69% | 68% | 63% | 70% |
| Sal_KS04_O | 73% | 79% | 74% | 74% | 74% | 70% | 67% | 73% | 74% | 70% | 76% | 68% | 69% | 72% | 69% | 68% | 66% | 70% | 68% | 64% | 69% |
| Sal_KS05_A | 70% | 73% | 73% | 73% | 72% | 72% | 69% | 70% | 71% | 71% | 73% | 69% | 68% | 70% | 67% | 69% | 66% | 70% | 70% | 63% | 68% |
| Sal_KS06_K | 72% | 77% | 74% | 75% | 72% | 74% | 71% | 72% | 76% | 71% | 75% | 70% | 70% | 71% | 71% | 69% | 68% | 70% | 72% | 64% | 68% |
| Sal_KS07_H | 71% | 74% | 70% | 73% | 70% | 71% | 69% | 72% | 71% | 67% | 71% | 66% | 66% | 68% | 68% | 66% | 65% | 69% | 67% | 62% | 68% |
| Sal_KS08_K | 72% | 69% | 70% | 69% | 71% | 69% | 67% | 67% | 69% | 70% | 70% | 67% | 65% | 67% | 64% | 64% | 63% | 65% | 64% | 65% | 64% |
| Sal_KS09_A | 69% | 71% | 72% | 72% | 73% | 69% | 66% | 71% | 72% | 68% | 76% | 69% | 66% | 69% | 67% | 66% | 66% | 68% | 66% | 64% | 66% |
| Sal_KS10_H | 68% | 69% | 70% | 68% | 69% | 74% | 67% | 68% | 69% | 66% | 73% | 68% | 66% | 68% | 69% | 66% | 65% | 69% | 67% | 64% | 69% |
| Sal_KS11_K | 71% | 71% | 71% | 70% | 73% | 70% | 69% | 70% | 71% | 68% | 74% | 68% | 67% | 68% | 68% | 63% | 65% | 65% | 67% | 64% | 65% |
| Sal_KS12_H | 71% | 72% | 73% | 72% | 75% | 72% | 65% | 75% | 74% | 71% | 75% | 69% | 67% | 68% | 70% | 67% | 67% | 68% | 68% | 63% | 67% |
| Sal_KS13_H | 71% | 74% | 71% | 73% | 72% | 70% | 68% | 70% | 73% | 68% | 73% | 69% | 65% | 68% | 67% | 68% | 65% | 70% | 68% | 65% | 68% |
| Sal_KS14_A | 66% | 68% | 69% | 68% | 68% | 67% | 64% | 65% | 67% | 64% | 69% | 64% | 65% | 63% | 64% | 61% | 62% | 63% | 65% | 61% | 67% |

| | Mon_KS12_H | Mon_KS_01 | Nan_KS01 | Nan_KS02_K | Nan_KS03_A | Nan_KS04_O | Nan_KS05_A | Nan_KS06_K | Nan_KS07_H | Nan_KS08_O | Nan_KS09_A | Nan_KS10_K | Nan_KS11_H | Nan_KS12_H | Nig_KS01 | Nig_KS02_K | Nig_KS03_A | Nig_KS04_O | Nig_KS05_A | Nig_KS06_O | Nig_KS07_A | Nig_KS08_O |
|---|---|---|---|---|---|---|---|---|---|---|---|---|---|---|---|---|---|---|---|---|---|---|
| Lsd_KS01 | 69% | 74% | 73% | 72% | 70% | 74% | 70% | 68% | 68% | 69% | 70% | 67% | 73% | 74% | 75% | 71% | 71% | 74% | 72% | 71% | 71% | 72% |
| Lsd_KS02_O | 72% | 74% | 70% | 69% | 70% | 73% | 72% | 70% | 69% | 71% | 73% | 67% | 75% | 75% | 74% | 71% | 73% | 75% | 71% | 75% | 71% | 74% |
| Lsd_KS03_A | 67% | 71% | 72% | 70% | 69% | 71% | 71% | 70% | 68% | 72% | 69% | 68% | 74% | 74% | 73% | 70% | 71% | 72% | 73% | 73% | 72% | 73% |
| Lsd_KS04_O | 71% | 74% | 73% | 73% | 71% | 74% | 72% | 72% | 71% | 73% | 73% | 70% | 74% | 74% | 74% | 72% | 73% | 76% | 73% | 76% | 73% | 77% |
| Lsd_KS05_A | 69% | 73% | 74% | 70% | 70% | 74% | 72% | 70% | 66% | 72% | 70% | 67% | 73% | 74% | 73% | 70% | 70% | 74% | 72% | 74% | 70% | 74% |
| Lsd_KS06_H | 71% | 70% | 71% | 68% | 69% | 72% | 71% | 68% | 70% | 68% | 72% | 67% | 74% | 75% | 73% | 69% | 69% | 73% | 70% | 71% | 69% | 70% |
| Lsd_KS07_K | 70% | 65% | 67% | 68% | 65% | 67% | 69% | 65% | 65% | 65% | 69% | 64% | 69% | 71% | 68% | 69% | 65% | 68% | 69% | 69% | 67% | 68% |
| Lsd_KS08_H | 69% | 69% | 71% | 69% | 68% | 71% | 69% | 68% | 68% | 68% | 70% | 66% | 71% | 74% | 72% | 70% | 70% | 72% | 70% | 72% | 68% | 71% |
| Lsd_KS09_A | 68% | 71% | 73% | 71% | 70% | 73% | 71% | 71% | 67% | 72% | 71% | 66% | 73% | 73% | 72% | 72% | 72% | 73% | 72% | 73% | 72% | 74% |
| Lsd_KS10_K | 66% | 67% | 67% | 68% | 66% | 68% | 66% | 66% | 66% | 67% | 68% | 66% | 70% | 71% | 69% | 68% | 68% | 70% | 67% | 70% | 68% | 70% |
| Lsd_KS11_O | 70% | 72% | 76% | 70% | 70% | 73% | 71% | 72% | 68% | 72% | 70% | 66% | 75% | 76% | 73% | 73% | 73% | 73% | 73% | 75% | 73% | 74% |
| Mon_KS02_K | 69% | 70% | 70% | 75% | 68% | 70% | 69% | 68% | 68% | 69% | 75% | 66% | 71% | 71% | 72% | 78% | 68% | 71% | 71% | 72% | 71% | 71% |
| Mon_KS03_A | 69% | 72% | 71% | 67% | 81% | 70% | 72% | 70% | 70% | 71% | 70% | 66% | 68% | 69% | 72% | 69% | 77% | 71% | 75% | 72% | 73% | 72% |
| Mon_KS04_O | 73% | 75% | 73% | 70% | 72% | 81% | 73% | 72% | 67% | 71% | 75% | 67% | 71% | 71% | 76% | 73% | 72% | 81% | 72% | 73% | 72% | 72% |
| Mon_KS05_A | 69% | 72% | 72% | 69% | 71% | 70% | 77% | 71% | 70% | 71% | 73% | 67% | 70% | 71% | 71% | 70% | 74% | 71% | 78% | 72% | 75% | 71% |
| Mon_KS06_O | 66% | 70% | 71% | 67% | 69% | 68% | 70% | 73% | 65% | 73% | 71% | 64% | 69% | 70% | 71% | 69% | 71% | 70% | 70% | 78% | 69% | 78% |
| Mon_KS07_A | 68% | 72% | 69% | 69% | 71% | 68% | 71% | 70% | 72% | 69% | 71% | 63% | 68% | 69% | 71% | 71% | 75% | 71% | 73% | 72% | 78% | 73% |
| Mon_KS08_O | 69% | 72% | 72% | 68% | 70% | 70% | 70% | 76% | 67% | 76% | 73% | 66% | 71% | 72% | 74% | 70% | 73% | 73% | 72% | 82% | 71% | 84% |
| Mon_KS09_A | 71% | 70% | 71% | 73% | 70% | 71% | 73% | 71% | 71% | 74% | 79% | 66% | 71% | 71% | 72% | 76% | 71% | 73% | 75% | 74% | 71% | 72% |
| Mon_KS10_K | 69% | 66% | 65% | 64% | 65% | 64% | 65% | 65% | 61% | 64% | 65% | 72% | 64% | 66% | 68% | 69% | 65% | 66% | 65% | 66% | 68% | 66% |
| Mon_KS11_H | 100% | 71% | 72% | 67% | 71% | 71% | 69% | 69% | 67% | 68% | 70% | 68% | 74% | 77% | 72% | 73% | 69% | 73% | 70% | 72% | 70% | 71% |
| Mon_KS12_H | 100% | 71% | 72% | 67% | 71% | 71% | 69% | 69% | 67% | 68% | 70% | 68% | 74% | 77% | 72% | 73% | 69% | 73% | 70% | 72% | 70% | 71% |
| Mon_KS_01 | 72% | 100% | 77% | 69% | 71% | 74% | 72% | 70% | 68% | 71% | 73% | 65% | 72% | 72% | 79% | 72% | 74% | 75% | 73% | 73% | 72% | 73% |
| Nan_KS01 | 71% | 76% | 100% | 70% | 69% | 74% | 72% | 73% | 68% | 74% | 71% | 65% | 71% | 71% | 79% | 71% | 72% | 74% | 72% | 74% | 70% | 73% |
| Nan_KS02_K | 67% | 69% | 70% | 100% | 67% | 71% | 69% | 69% | 66% | 71% | 73% | 65% | 68% | 68% | 69% | 78% | 69% | 72% | 70% | 72% | 70% | 72% |
| Nan_KS03_A | 71% | 71% | 71% | 68% | 100% | 70% | 75% | 70% | 67% | 71% | 72% | 67% | 72% | 71% | 70% | 70% | 78% | 72% | 74% | 73% | 73% | 73% |
| Nan_KS04_O | 71% | 74% | 75% | 71% | 69% | 100% | 71% | 73% | 67% | 71% | 71% | 67% | 71% | 71% | 75% | 73% | 72% | 83% | 72% | 75% | 70% | 73% |
| Nan_KS05_A | 69% | 72% | 73% | 69% | 75% | 71% | 100% | 71% | 70% | 72% | 79% | 67% | 70% | 71% | 72% | 72% | 76% | 72% | 80% | 73% | 73% | 72% |
| Nan_KS06_K | 69% | 70% | 74% | 69% | 70% | 73% | 71% | 100% | 66% | 85% | 72% | 66% | 71% | 72% | 72% | 71% | 71% | 73% | 69% | 79% | 71% | 79% |
| Nan_KS07_H | 67% | 68% | 69% | 66% | 67% | 68% | 70% | 67% | 100% | 68% | 69% | 63% | 67% | 68% | 71% | 67% | 70% | 70% | 71% | 72% | 73% | 71% |
| Nan_KS08_O | 68% | 71% | 75% | 71% | 71% | 71% | 71% | 85% | 68% | 100% | 74% | 66% | 71% | 70% | 73% | 73% | 73% | 73% | 71% | 80% | 72% | 81% |
| Nan_KS09_A | 71% | 73% | 72% | 73% | 72% | 72% | 79% | 72% | 69% | 74% | 100% | 65% | 71% | 72% | 73% | 77% | 72% | 74% | 74% | 74% | 71% | 73% |
| Nan_KS10_K | 68% | 65% | 66% | 65% | 66% | 67% | 67% | 66% | 63% | 66% | 65% | 100% | 67% | 68% | 68% | 67% | 67% | 69% | 67% | 68% | 67% | 69% |
| Nan_KS11_H | 74% | 72% | 72% | 68% | 72% | 71% | 69% | 71% | 67% | 71% | 71% | 67% | 100% | 84% | 73% | 69% | 71% | 74% | 72% | 72% | 71% | 73% |
| Nan_KS12_H | 77% | 71% | 72% | 68% | 71% | 71% | 71% | 72% | 68% | 70% | 72% | 68% | 84% | 100% | 74% | 70% | 70% | 73% | 72% | 73% | 70% | 74% |
| Nig_KS01 | 72% | 79% | 80% | 69% | 70% | 75% | 72% | 72% | 71% | 73% | 73% | 68% | 73% | 74% | 100% | 73% | 75% | 80% | 74% | 76% | 74% | 76% |
| Nig_KS02_K | 73% | 72% | 72% | 78% | 70% | 73% | 72% | 72% | 67% | 73% | 77% | 67% | 70% | 70% | 73% | 100% | 74% | 74% | 74% | 74% | 76% | 73% |
| Nig_KS03_A | 70% | 74% | 74% | 70% | 78% | 73% | 76% | 71% | 70% | 74% | 72% | 68% | 71% | 70% | 76% | 74% | 100% | 75% | 79% | 75% | 77% | 75% |
| Nig_KS04_O | 73% | 74% | 75% | 72% | 72% | 83% | 72% | 73% | 69% | 73% | 73% | 69% | 74% | 73% | 80% | 74% | 74% | 100% | 75% | 78% | 74% | 77% |
| Nig_KS05_A | 71% | 73% | 74% | 71% | 74% | 73% | 80% | 70% | 71% | 71% | 74% | 67% | 73% | 73% | 74% | 75% | 79% | 76% | 100% | 78% | 74% | 76% |
| Nig_KS06_O | 72% | 73% | 75% | 72% | 73% | 75% | 72% | 79% | 72% | 80% | 74% | 68% | 72% | 73% | 76% | 74% | 75% | 78% | 77% | 100% | 75% | 93% |
| Nig_KS07_A | 71% | 72% | 71% | 71% | 73% | 71% | 73% | 72% | 73% | 73% | 72% | 68% | 72% | 71% | 75% | 76% | 77% | 75% | 75% | 76% | 100% | 75% |
| Nig_KS08_O | 71% | 73% | 74% | 72% | 73% | 73% | 71% | 79% | 71% | 81% | 73% | 69% | 73% | 74% | 76% | 73% | 75% | 77% | 76% | 93% | 75% | 100% |
| Nig_KS09_A | 72% | 73% | 72% | 75% | 71% | 72% | 72% | 69% | 69% | 73% | 76% | 68% | 72% | 71% | 73% | 81% | 74% | 76% | 76% | 75% | 75% | 74% |
| Nig_KS10_K | 67% | 65% | 67% | 66% | 67% | 68% | 66% | 68% | 66% | 68% | 66% | 74% | 67% | 69% | 69% | 68% | 67% | 70% | 70% | 72% | 68% | 72% |
| Nig_KS11_H | 74% | 72% | 73% | 67% | 71% | 75% | 71% | 72% | 69% | 72% | 70% | 67% | 73% | 73% | 78% | 70% | 72% | 79% | 74% | 75% | 72% | 75% |
| Nig_KS12_H | 75% | 72% | 72% | 68% | 71% | 73% | 71% | 70% | 67% | 71% | 72% | 69% | 75% | 76% | 75% | 72% | 70% | 77% | 72% | 74% | 70% | 75% |
| Nig_KS13_H | 75% | 71% | 71% | 67% | 68% | 71% | 71% | 69% | 66% | 69% | 71% | 65% | 72% | 74% | 73% | 71% | 69% | 74% | 72% | 72% | 69% | 72% |
| Nig_KS14_A | 74% | 71% | 72% | 70% | 69% | 72% | 71% | 69% | 68% | 70% | 69% | 69% | 73% | 74% | 75% | 73% | 71% | 75% | 70% | 73% | 73% | 72% |
| Sal_KS01 | 67% | 70% | 71% | 66% | 66% | 69% | 67% | 66% | 65% | 66% | 67% | 64% | 67% | 69% | 70% | 68% | 67% | 70% | 69% | 70% | 68% | 68% |
| Sal_KS02-O | 72% | 73% | 73% | 66% | 69% | 70% | 68% | 69% | 66% | 69% | 70% | 63% | 72% | 73% | 73% | 68% | 70% | 72% | 70% | 73% | 69% | 72% |
| Sal_KS03_A | 70% | 70% | 70% | 68% | 69% | 69% | 69% | 65% | 66% | 67% | 69% | 64% | 71% | 74% | 69% | 67% | 69% | 69% | 69% | 71% | 70% | 70% |
| Sal_KS04_O | 69% | 72% | 71% | 65% | 69% | 72% | 69% | 70% | 68% | 70% | 67% | 66% | 73% | 72% | 72% | 67% | 69% | 73% | 69% | 74% | 70% | 73% |
| Sal_KS05_A | 68% | 70% | 68% | 68% | 69% | 70% | 68% | 68% | 66% | 67% | 68% | 65% | 73% | 73% | 70% | 70% | 70% | 72% | 70% | 70% | 70% | 72% |
| Sal_KS06_K | 68% | 69% | 70% | 69% | 71% | 70% | 73% | 70% | 69% | 69% | 72% | 66% | 72% | 73% | 72% | 71% | 71% | 72% | 73% | 74% | 73% | 73% |
| Sal_KS07_H | 68% | 69% | 69% | 66% | 65% | 68% | 69% | 65% | 65% | 65% | 67% | 63% | 69% | 72% | 72% | 66% | 68% | 72% | 68% | 71% | 68% | 69% |
| Sal_KS08_K | 64% | 66% | 66% | 66% | 64% | 69% | 65% | 66% | 62% | 64% | 65% | 64% | 67% | 69% | 68% | 65% | 65% | 68% | 64% | 66% | 65% | 66% |
| Sal_KS09_A | 66% | 67% | 68% | 67% | 66% | 70% | 66% | 65% | 65% | 65% | 66% | 64% | 68% | 70% | 70% | 69% | 68% | 70% | 68% | 69% | 67% | 68% |
| Sal_KS10_H | 69% | 66% | 68% | 64% | 66% | 69% | 68% | 69% | 64% | 67% | 69% | 65% | 69% | 71% | 68% | 66% | 67% | 69% | 67% | 69% | 66% | 70% |
| Sal_KS11_K | 65% | 67% | 66% | 65% | 66% | 68% | 66% | 68% | 64% | 66% | 69% | 63% | 68% | 70% | 68% | 68% | 65% | 68% | 68% | 69% | 67% | 68% |
| Sal_KS12_H | 67% | 70% | 70% | 67% | 67% | 69% | 68% | 68% | 65% | 68% | 69% | 66% | 72% | 72% | 71% | 68% | 68% | 70% | 68% | 69% | 68% | 69% |
| Sal_KS13_H | 68% | 71% | 69% | 67% | 67% | 70% | 69% | 67% | 67% | 68% | 68% | 64% | 70% | 71% | 71% | 67% | 68% | 71% | 70% | 74% | 67% | 72% |
| Sal_KS14_A | 67% | 66% | 70% | 65% | 65% | 65% | 66% | 64% | 63% | 67% | 64% | 61% | 66% | 66% | 70% | 67% | 65% | 68% | 65% | 67% | 65% | 66% |

| | Nig_KS09_A | Nig_KS10_K | Nig_KS11_H | Nig_KS12_H | Nig_KS13_H | Nig_KS14_A | Sal_KS01 | Sal_KS02-O | Sal_KS03_A | Sal_KS04_O | Sal_KS05_A | Sal_KS06_K | Sal_KS07_H | Sal_KS08_K | Sal_KS09_A | Sal_KS10_H | Sal_KS11_K | Sal_KS12_H | Sal_KS13_H | Sal_KS14_A |
|---|---|---|---|---|---|---|---|---|---|---|---|---|---|---|---|---|---|---|---|---|
| Lsd_KS01 | 71% | 68% | 72% | 72% | 70% | 71% | 76% | 71% | 71% | 73% | 71% | 72% | 71% | 72% | 69% | 69% | 71% | 71% | 71% | 66% |
| Lsd_KS02_O | 74% | 70% | 75% | 75% | 72% | 73% | 72% | 80% | 75% | 79% | 74% | 77% | 74% | 69% | 71% | 69% | 71% | 73% | 74% | 68% |
| Lsd_KS03_A | 73% | 69% | 71% | 74% | 69% | 73% | 69% | 71% | 75% | 74% | 73% | 74% | 70% | 70% | 72% | 70% | 71% | 73% | 71% | 68% |
| Lsd_KS04_O | 75% | 71% | 74% | 75% | 74% | 74% | 68% | 74% | 73% | 75% | 74% | 76% | 74% | 70% | 73% | 69% | 70% | 72% | 74% | 68% |
| Lsd_KS05_A | 71% | 70% | 72% | 73% | 70% | 73% | 71% | 73% | 75% | 75% | 73% | 73% | 71% | 72% | 73% | 70% | 73% | 75% | 72% | 68% |
| Lsd_KS06_H | 72% | 70% | 73% | 74% | 73% | 71% | 67% | 69% | 73% | 70% | 72% | 74% | 71% | 69% | 69% | 74% | 70% | 71% | 70% | 66% |
| Lsd_KS07_K | 70% | 68% | 68% | 69% | 68% | 67% | 68% | 69% | 69% | 67% | 69% | 71% | 69% | 66% | 66% | 67% | 69% | 65% | 68% | 64% |
| Lsd_KS08_H | 72% | 69% | 72% | 71% | 68% | 72% | 67% | 72% | 72% | 73% | 70% | 72% | 72% | 67% | 71% | 68% | 70% | 75% | 70% | 65% |
| Lsd_KS09_A | 72% | 70% | 72% | 72% | 70% | 73% | 72% | 73% | 73% | 74% | 71% | 76% | 72% | 69% | 72% | 70% | 71% | 74% | 73% | 67% |
| Lsd_KS10_K | 68% | 68% | 70% | 69% | 66% | 68% | 67% | 69% | 68% | 70% | 71% | 71% | 67% | 70% | 68% | 66% | 68% | 71% | 68% | 63% |
| Lsd_KS11_O | 73% | 71% | 72% | 74% | 72% | 75% | 72% | 74% | 74% | 76% | 73% | 75% | 71% | 70% | 75% | 73% | 73% | 75% | 73% | 68% |
| Mon_KS02_K | 76% | 67% | 69% | 70% | 69% | 70% | 68% | 69% | 67% | 68% | 70% | 70% | 67% | 67% | 69% | 68% | 68% | 69% | 69% | 64% |
| Mon_KS03_A | 73% | 65% | 72% | 68% | 68% | 70% | 68% | 68% | 67% | 69% | 68% | 70% | 66% | 65% | 66% | 66% | 67% | 67% | 65% | 65% |
| Mon_KS04_O | 76% | 66% | 74% | 73% | 72% | 72% | 71% | 70% | 67% | 72% | 70% | 71% | 68% | 67% | 69% | 68% | 67% | 68% | 67% | 63% |
| Mon_KS05_A | 73% | 66% | 70% | 68% | 69% | 71% | 71% | 68% | 68% | 70% | 67% | 71% | 68% | 64% | 67% | 69% | 68% | 71% | 67% | 64% |
| Mon_KS06_O | 70% | 68% | 69% | 69% | 69% | 68% | 65% | 67% | 67% | 68% | 69% | 69% | 66% | 64% | 66% | 66% | 63% | 67% | 68% | 60% |
| Mon_KS07_A | 73% | 66% | 69% | 68% | 67% | 67% | 68% | 67% | 67% | 67% | 67% | 69% | 66% | 63% | 67% | 65% | 65% | 67% | 65% | 62% |
| Mon_KS08_O | 71% | 68% | 72% | 71% | 71% | 69% | 66% | 70% | 68% | 71% | 71% | 71% | 69% | 65% | 68% | 69% | 65% | 68% | 70% | 63% |
| Mon_KS09_A | 80% | 68% | 73% | 72% | 69% | 70% | 68% | 69% | 68% | 68% | 71% | 72% | 67% | 64% | 66% | 67% | 67% | 68% | 68% | 64% |
| Mon_KS10_K | 67% | 72% | 64% | 67% | 65% | 66% | 62% | 64% | 63% | 64% | 64% | 64% | 62% | 65% | 64% | 64% | 64% | 63% | 64% | 61% |
| Mon_KS11_H | 73% | 67% | 74% | 75% | 75% | 74% | 67% | 72% | 69% | 69% | 68% | 68% | 68% | 64% | 66% | 69% | 65% | 67% | 68% | 66% |
| Mon_KS12_H | 73% | 67% | 74% | 75% | 75% | 74% | 67% | 72% | 69% | 69% | 68% | 68% | 68% | 64% | 66% | 69% | 65% | 67% | 68% | 66% |
| Mon_KS_01 | 74% | 65% | 72% | 72% | 71% | 72% | 71% | 73% | 70% | 72% | 70% | 70% | 69% | 66% | 67% | 67% | 67% | 70% | 71% | 66% |
| Nan_KS01 | 72% | 66% | 72% | 71% | 70% | 71% | 70% | 72% | 69% | 70% | 67% | 69% | 68% | 65% | 68% | 67% | 65% | 69% | 68% | 69% |
| Nan_KS02_K | 76% | 66% | 67% | 68% | 67% | 70% | 66% | 66% | 68% | 65% | 68% | 69% | 66% | 66% | 67% | 64% | 65% | 67% | 66% | 65% |
| Nan_KS03_A | 72% | 67% | 71% | 71% | 68% | 70% | 67% | 69% | 69% | 69% | 69% | 71% | 66% | 64% | 67% | 67% | 66% | 67% | 67% | 65% |
| Nan_KS04_O | 73% | 68% | 74% | 73% | 71% | 72% | 69% | 70% | 68% | 72% | 70% | 70% | 68% | 69% | 70% | 69% | 67% | 69% | 69% | 64% |
| Nan_KS05_A | 73% | 67% | 71% | 71% | 71% | 71% | 67% | 69% | 69% | 69% | 68% | 73% | 70% | 65% | 66% | 68% | 66% | 69% | 69% | 66% |
| Nan_KS06_K | 70% | 68% | 72% | 70% | 69% | 69% | 66% | 69% | 65% | 70% | 68% | 70% | 65% | 66% | 65% | 69% | 67% | 68% | 66% | 64% |
| Nan_KS07_H | 70% | 67% | 70% | 68% | 66% | 68% | 65% | 67% | 66% | 68% | 67% | 69% | 65% | 62% | 65% | 65% | 64% | 65% | 67% | 63% |
| Nan_KS08_O | 74% | 68% | 72% | 71% | 69% | 70% | 66% | 69% | 67% | 70% | 67% | 69% | 65% | 64% | 65% | 67% | 66% | 68% | 68% | 66% |
| Nan_KS09_A | 78% | 66% | 71% | 72% | 71% | 69% | 67% | 70% | 68% | 68% | 69% | 72% | 68% | 65% | 66% | 70% | 69% | 69% | 68% | 64% |
| Nan_KS10_K | 69% | 74% | 67% | 69% | 65% | 69% | 64% | 63% | 63% | 66% | 65% | 66% | 63% | 64% | 64% | 65% | 63% | 66% | 64% | 61% |
| Nan_KS11_H | 73% | 67% | 73% | 75% | 72% | 73% | 67% | 72% | 71% | 73% | 73% | 72% | 69% | 66% | 68% | 69% | 68% | 72% | 70% | 66% |
| Nan_KS12_H | 72% | 69% | 73% | 76% | 74% | 74% | 69% | 73% | 73% | 72% | 73% | 73% | 72% | 69% | 69% | 71% | 69% | 72% | 71% | 66% |
| Nig_KS01 | 74% | 69% | 78% | 75% | 73% | 75% | 70% | 73% | 69% | 72% | 70% | 72% | 72% | 68% | 70% | 68% | 68% | 71% | 71% | 69% |
| Nig_KS02_K | 82% | 68% | 70% | 72% | 71% | 73% | 68% | 68% | 67% | 67% | 70% | 71% | 66% | 65% | 69% | 66% | 68% | 68% | 67% | 67% |
| Nig_KS03_A | 76% | 68% | 72% | 71% | 70% | 71% | 68% | 70% | 69% | 70% | 70% | 72% | 68% | 65% | 69% | 68% | 65% | 68% | 68% | 65% |
| Nig_KS04_O | 77% | 70% | 78% | 77% | 74% | 75% | 70% | 72% | 69% | 73% | 72% | 72% | 71% | 67% | 69% | 69% | 67% | 70% | 70% | 67% |
| Nig_KS05_A | 78% | 70% | 74% | 72% | 73% | 71% | 69% | 71% | 69% | 69% | 71% | 73% | 68% | 64% | 68% | 67% | 68% | 69% | 70% | 65% |
| Nig_KS06_O | 76% | 72% | 75% | 74% | 72% | 73% | 70% | 73% | 71% | 74% | 71% | 74% | 71% | 66% | 69% | 69% | 69% | 69% | 73% | 67% |
| Nig_KS07_A | 76% | 69% | 73% | 71% | 69% | 74% | 68% | 70% | 71% | 70% | 71% | 74% | 69% | 66% | 67% | 67% | 68% | 68% | 67% | 65% |
| Nig_KS08_O | 75% | 72% | 75% | 75% | 72% | 72% | 68% | 72% | 70% | 73% | 72% | 73% | 69% | 66% | 68% | 70% | 68% | 69% | 71% | 66% |
| Nig_KS09_A | 100% | 69% | 71% | 74% | 70% | 74% | 70% | 72% | 68% | 70% | 73% | 73% | 69% | 65% | 69% | 67% | 68% | 69% | 70% | 66% |
| Nig_KS10_K | 70% | 100% | 74% | 72% | 69% | 70% | 64% | 67% | 67% | 67% | 68% | 69% | 67% | 66% | 66% | 65% | 66% | 66% | 68% | 64% |
| Nig_KS11_H | 72% | 74% | 100% | 82% | 74% | 75% | 69% | 74% | 69% | 73% | 72% | 74% | 73% | 67% | 68% | 69% | 66% | 69% | 70% | 68% |
| Nig_KS12_H | 75% | 72% | 82% | 100% | 77% | 79% | 68% | 74% | 70% | 72% | 72% | 73% | 69% | 68% | 69% | 69% | 66% | 69% | 70% | 68% |
| Nig_KS13_H | 71% | 69% | 74% | 77% | 100% | 75% | 65% | 70% | 67% | 71% | 68% | 69% | 66% | 64% | 65% | 70% | 65% | 66% | 67% | 64% |
| Nig_KS14_A | 75% | 70% | 75% | 79% | 75% | 100% | 69% | 72% | 69% | 72% | 68% | 71% | 70% | 65% | 67% | 68% | 66% | 69% | 68% | 65% |
| Sal_KS01 | 71% | 64% | 69% | 68% | 65% | 69% | 100% | 70% | 67% | 71% | 67% | 71% | 66% | 67% | 71% | 67% | 70% | 67% | 70% | 67% |
| Sal_KS02-O | 73% | 67% | 74% | 74% | 70% | 72% | 70% | 100% | 71% | 77% | 70% | 74% | 70% | 68% | 69% | 67% | 70% | 69% | 74% | 67% |
| Sal_KS03_A | 69% | 67% | 70% | 70% | 68% | 70% | 67% | 72% | 100% | 72% | 72% | 72% | 71% | 65% | 69% | 68% | 68% | 71% | 71% | 66% |
| Sal_KS04_O | 71% | 67% | 73% | 72% | 71% | 72% | 71% | 77% | 72% | 100% | 71% | 76% | 71% | 68% | 69% | 69% | 70% | 71% | 72% | 66% |
| Sal_KS05_A | 74% | 67% | 71% | 72% | 68% | 68% | 67% | 70% | 72% | 71% | 100% | 77% | 70% | 69% | 70% | 67% | 67% | 69% | 71% | 65% |
| Sal_KS06_K | 74% | 69% | 74% | 73% | 69% | 71% | 71% | 74% | 72% | 76% | 77% | 100% | 71% | 70% | 71% | 69% | 72% | 71% | 74% | 68% |
| Sal_KS07_H | 69% | 67% | 73% | 69% | 66% | 70% | 66% | 70% | 71% | 71% | 71% | 71% | 100% | 66% | 67% | 65% | 65% | 68% | 72% | 65% |
| Sal_KS08_K | 66% | 66% | 67% | 68% | 64% | 65% | 68% | 68% | 65% | 68% | 69% | 70% | 66% | 100% | 71% | 66% | 67% | 69% | 69% | 65% |
| Sal_KS09_A | 70% | 67% | 68% | 69% | 65% | 67% | 71% | 69% | 68% | 69% | 71% | 71% | 67% | 71% | 100% | 67% | 70% | 69% | 70% | 66% |
| Sal_KS10_H | 68% | 65% | 69% | 69% | 70% | 68% | 67% | 67% | 68% | 69% | 67% | 69% | 65% | 66% | 67% | 100% | 66% | 68% | 66% | 63% |
| Sal_KS11_K | 69% | 66% | 66% | 66% | 65% | 66% | 70% | 70% | 68% | 71% | 68% | 72% | 65% | 67% | 70% | 67% | 100% | 71% | 69% | 64% |
| Sal_KS12_H | 70% | 67% | 69% | 69% | 66% | 70% | 67% | 70% | 70% | 72% | 70% | 71% | 68% | 69% | 69% | 68% | 71% | 100% | 69% | 65% |
| Sal_KS13_H | 71% | 68% | 70% | 71% | 67% | 68% | 70% | 75% | 71% | 73% | 71% | 74% | 72% | 69% | 71% | 67% | 70% | 70% | 100% | 68% |
| Sal_KS14_A | 67% | 64% | 68% | 68% | 64% | 66% | 67% | 67% | 66% | 66% | 65% | 68% | 65% | 65% | 66% | 63% | 64% | 66% | 68% | 100% |

Figure S 1 Distance matrix generated with the “Simple similarity” algorithm in UGENE v48.1 to identify evolutionarily related polyether producers. Four producers with 61–100% sequence identity were selected for phylogenetic analysis, with NCBI GenBank IDs as follows: Nig (*DQ354110.1*), Mon (*AF440781.1*), Nan (*AF521085.1*), Sal (*JN033543.1*), and Tmn (*AB193609.1*).

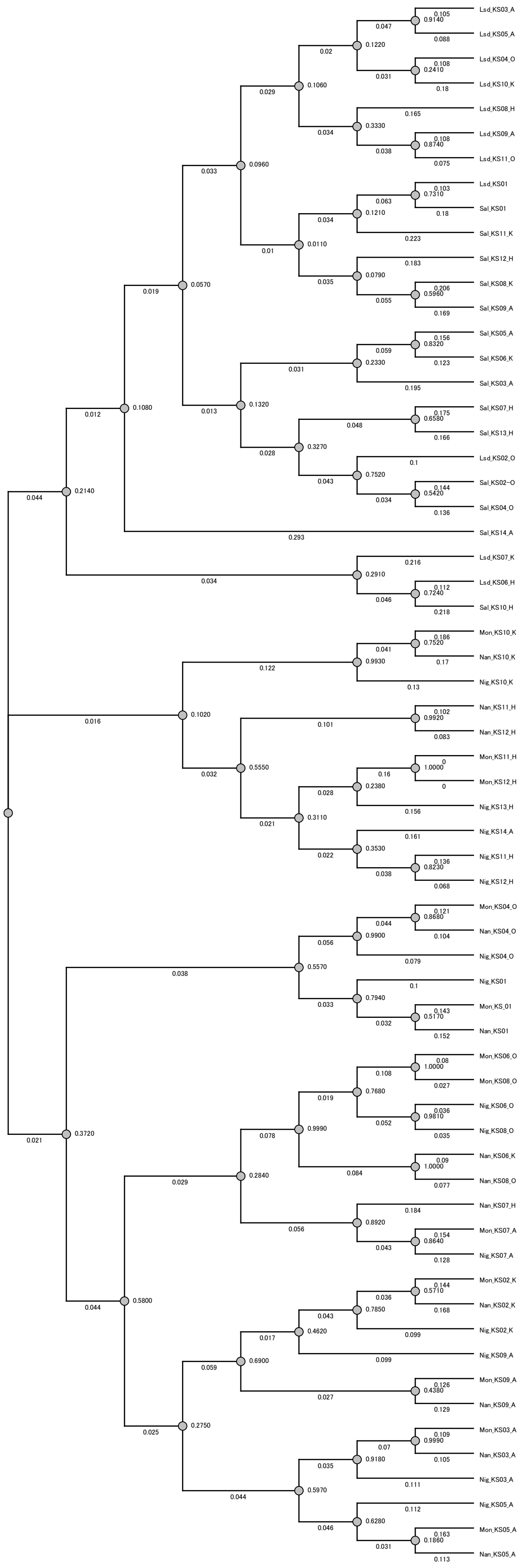


Figure S 2 Phylogenetic tree of KS domains with bootstrap value (knodes) and distance. Phylogenetic analyses were conducted using MEGA 7. Multiple sequence alignment was performed with ClustalW using default parameters. The evolutionary history was inferred using the Maximum Likelihood method with the Jones–Taylor–Thornton (JTT) model of amino acid substitution. Rates among sites were assumed to be uniform. Gaps and missing data were handled by partial deletion with a site coverage cutoff of 30%. Initial trees for the Maximum Likelihood analysis were generated automatically by Neighbor-Joining and BioNJ algorithms, with branch optimization performed using the Nearest-Neighbor-Interchange (NNI) heuristic search. Statistical support for the inferred tree topology was evaluated with 1000 bootstrap replicates.

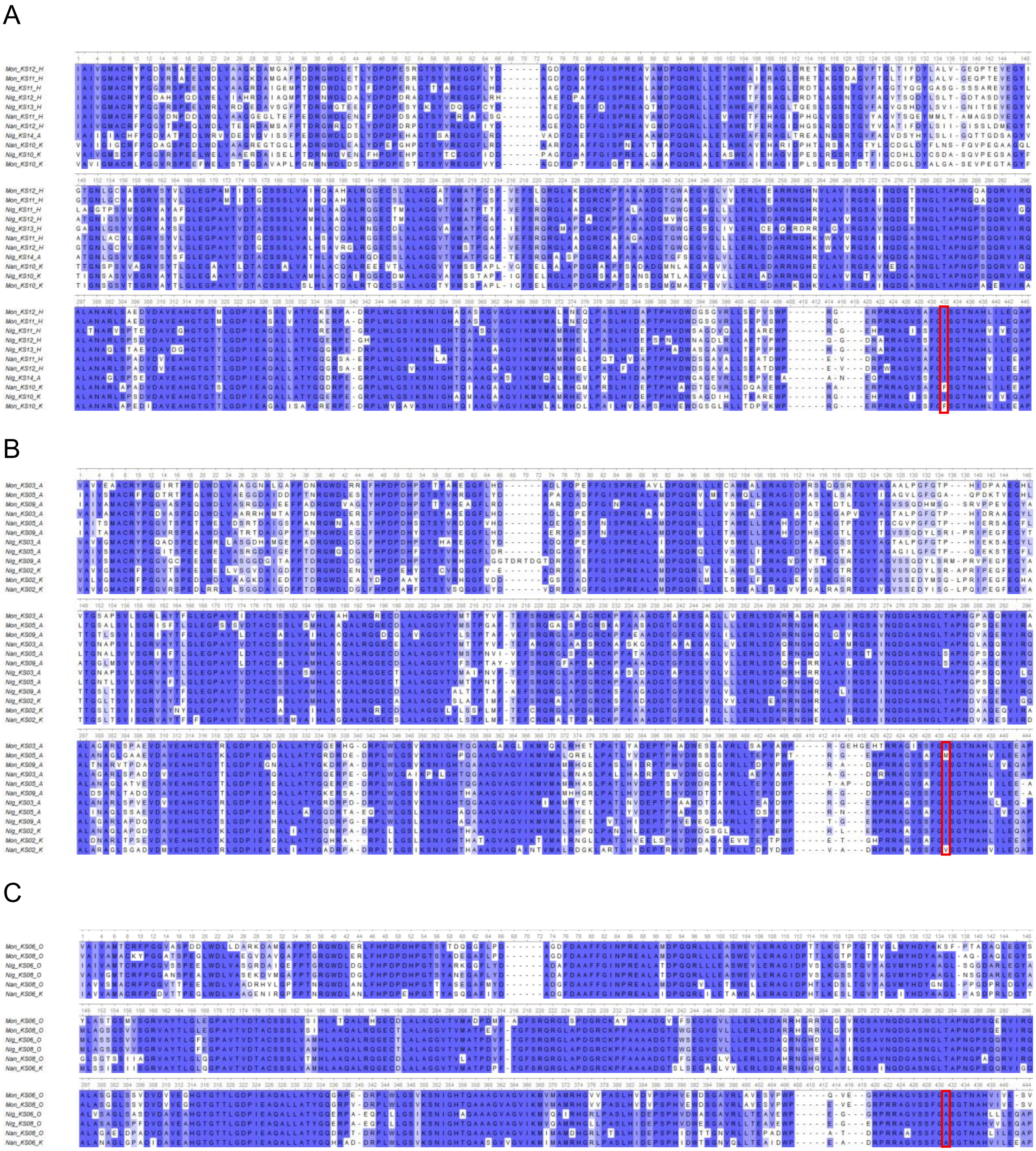

Figure S 3: Multiple sequence alignments of (A) H-clade (β-hydroxy), (B) A-clade (β-aliphatic), and (C) O-clade (β-olefinic) were performed using ClustalW with default settings in UGENE v48.1. Red boxes highlight motif V residue. Conservation level over 50% were highlighted in percentage identity color.

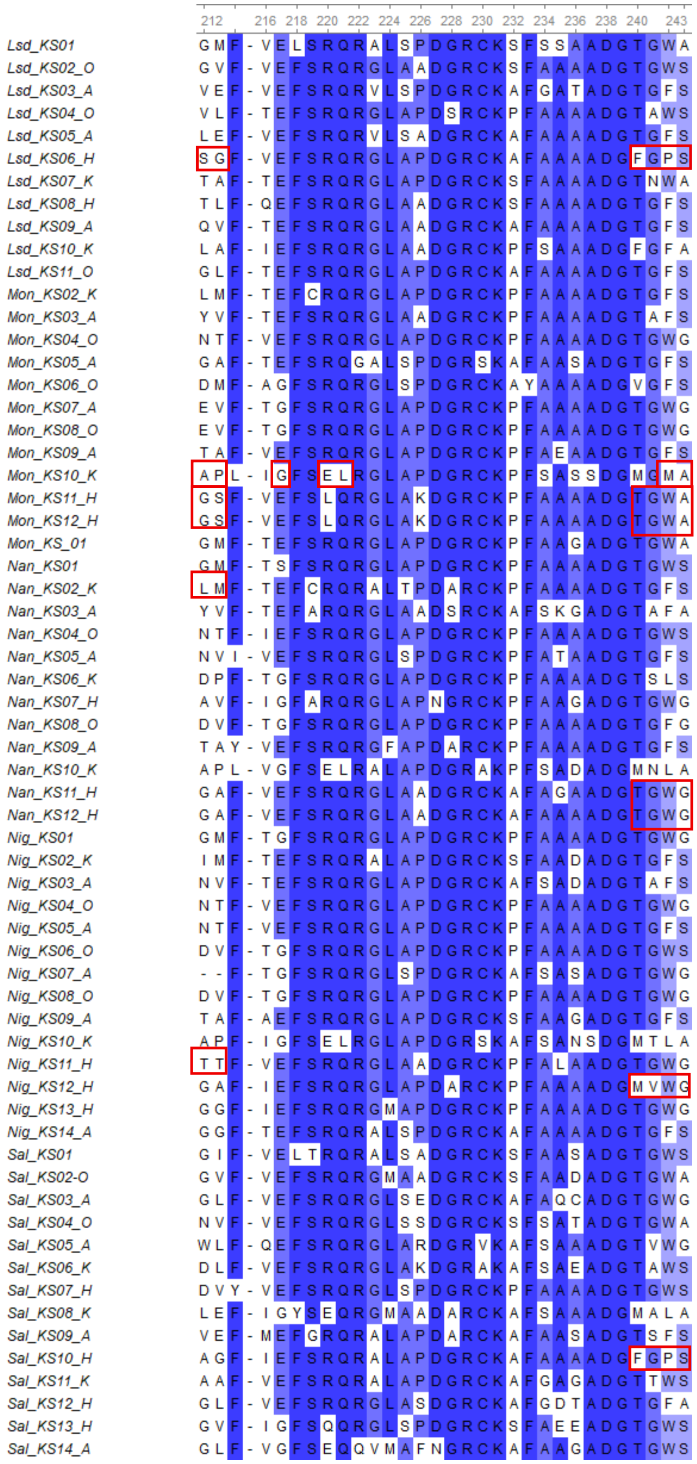


Figure S 4: Parts of multiple sequence alignments of all selected KS domains. MSA was generated with ClustalW (default settings) in UGENE v48.1. Residues with >50% conservation are highlighted according to percentage identity coloring. Red boxes highlight the template residues that were used for mutation design. Numbers above the sequence represent the column number of the alignment.

### B) Materials and methods

#### KS5 mutagenesis

A KS5 deletion construct was generated by replacing KS5 with a spectinomycin resistance ($Spt^R$) cassette. Overlap extension PCR fragments (primers 344–1978, 1977–345, 723–724) were cloned into HindIII/EcoRI-digested pKC1139, a vector with a temperature-sensitive replicon and apramycin resistance ($Apr^R$), using the GC-SLIC method [1]. For counter-selection, *codA* from *E. coli* was inserted [2], and plasmids carrying the mutations were generated similarly with mutagenic primers 1963 and 1964. Constructs were introduced into *E. coli* ET12567/pUZ8002 and transferred into *Streptomyces cinnamonensis* A495 by conjugation. Exconjugants were first grown at 39 °C and then shifted to 30 °C to allow plasmid excision by a second crossover. Counter-selection with 5-fluorocytosine enriched for double recombinants, producing $Apr^S$ $Spt^R$ colonies used as recipients for subsequent conjugations. Mutant plasmids were introduced into this strain, and $Apr^S$ $Spt^S$ variants were selected. As a control, the wild-type KS5 gene (primers 344–345) was cloned and reintroduced into the $Apr^s$ $Spt^R$ strain, and final colonies were confirmed by colony PCR (primers 1981–1982). Oligonucleotides are listed in Table S1.

Table S 1 List of oligonucleotides used in this study.

| **Oligo number** | **Oligo name** | **Oligo sequence** |
|---|---|---|
| **344** | 344_KS5mut-1_for | atgacatgattacgaattctttgccgcggccgatgtcagtgatccggacgc |
| **345** | 345_KS5mut-2_rev | acggccagtgccaagctgtggtcgcggcgcagggtgggcacgaccgtgg |
| **723** | 723_res-marker_for | tgcagctcacggtaactgatgccgtatttgcagtaccagcg |
| **724** | 724_res-marker_rev | tggagctgcttcgaagttcctatactttctagagaataggaacttcgg |
| **1977** | 1977_KS5-Swa_for | tggccatggacccgcagcagcgggtgctcatttaaatcaagatggtgatggcgctgcagcgcg |
| **1978** | 1978_KS5-Swa_rev | ttcgcgctgcagcgccatcaccatcttgatttaaatgagcacccgctgctgcgggtccatggcc |
| **1981** | 1981_KS5-seq1_rev | ttcctcgccccggccggagacgacccacgggacgactgcg |
| **1982** | 1982_KS5-seq2_for | aagcgggtcacggtggacctcggccaggcccgccggcgg |
| **1913** | 1913_KS5-G214-L_for | tgtcaccgtgatgtcgacgcccctcgcgttcaccgagttctcccgccagggcgcgctg |
| **1914** | 1914_KS5-G214-L_rev | tggcgggagaactcggtgaacgcgaggggcgtcgacatcacggtgacaccgccggccagcg |
| **1915** | 1915_KS5-A215-M_for | tgtcaccgtgatgtcgacgcccggcatgttcaccgagttctcccgccagggcgcgctg |
| **1916** | 1916_KS5-A215-M_rev | tggcgggagaactcggtgaacatgccgggcgtcgacatcacggtgacaccgccggccagcg |
| **1917** | 1917_KS5-GA214_215-LM_for | tgtcaccgtgatgtcgacgcccctcatgttcaccgagttctcccgccagggcgcgctg |
| **1918** | 1918_KS5-GA214_215-LM_rev | tggcgggagaactcggtgaacatgaggggcgtcgacatcacggtgacaccgccggccagcg |
| **1919** | 1919_KS5-G214-A_for | tgtcaccgtgatgtcgacgcccgccgcgttcaccgagttctcccgccagggcgcgctg |
| **1920** | 1920_KS5-G214-A_rev | tggcgggagaactcggtgaacgcggcgggcgtcgacatcacggtgacaccgccggccagcg |
| **1921** | 1921_KS5-A215-P_for | tgtcaccgtgatgtcgacgcccggcccgttcaccgagttctcccgccagggcgcgctg |
| **1922** | 1922_KS5-A215-P_rev | tggcgggagaactcggtgaacgggccgggcgtcgacatcacggtgacaccgccggccagcg |
| **1923** | 1923_KS5-GA214_215-AP_for | tgtcaccgtgatgtcgacgcccgccccgttcaccgagttctcccgccagggcgcgctg |
| **1924** | 1924_KS5-GA214_215-AP_rev | tggcgggagaactcggtgaacggggcgggcgtcgacatcacggtgacaccgccggccagcg |

| | | |
|---|---|---|
| **1925** | 1925_KS5-A215-P_E218-G_for | tgtcaccgtgatgtcgacgcccggcccgttcaccgggttctcccgccagggcgcgctgtctccggacgg |
| **1926** | 1926_KS5-A215-P_E218-G_rev | acagcgcgccctggcgggagaacccggtgaacgggccgggcgtcgacatcacggtgacatcacggtgacaccg |
| **1927** | 1927_KS5-GA214_215-AP_E218-G_for | tgtcaccgtgatgtcgacgcccgccccgttcaccgggttctcccgccagggcgcgctgtctccggacgg |
| **1928** | 1928_KS5-GA214_215-AP_E218-G_rev | agacagcgcgccctggcgggagaacccggtgaacggggcgggcgtcgacatcacggtgacaccgccggc |
| **1929** | 1929_KS5-G214-A_E218-G_for | tgtcaccgtgatgtcgacgcccgccgcgttcaccgggttctcccgccagggcgcgctgtctccggacgg |
| **1930** | 1930_KS5-G214-A_E218-G_rev | agacagcgcgccctggcgggagaacccggtgaacgcggcgggcgtcgacatcacggtgacaccgccgg |
| **1931** | 1931_KS5-E218-G_for | tgtcaccgtgatgtcgacgcccggcgcgttcaccgggttctcccgccagggcgcgctgtctccggacgg |
| **1932** | 1932_KS5-E218-G_rev | tccggagacagcgcgccctggcgggagaacccggtgaacgcgccgggcgtcgacatcacgg |
| **1933** | 1933_KS5-R221-E_for | acgcccggcgcgttcaccgagttctccgagcagggcgcgctgtctccggacggccgctcg |
| **1934** | 1934_KS5-R221-E_rev | ttcgagcggccgtccggagacagcgcgccctgctcggagaactcggtgaacgcgccgggcg |
| **1935** | 1935_KS5-Q222-L_for | acgcccggcgcgttcaccgagttctcccgcctgggcgcgctgtctccggacggccgctcg |
| **1936** | 1936_KS5-Q222-L_rev | ttcgagcggccgtccggagacagcgcgcccaggcgggagaactcggtgaacgcgccgggcg |
| **1937** | 1937_KS5-F243-M_for | ttcgcggcctcggccgacggcaccggtatgtcggagggcgcgggactgctcctcctggagcgg |
| **1938** | 1938_KS5-F243-M_rev | aggagcagtcccgcgccctccgacataccggtgccgtcggccgaggccgcgaaagccttcg |
| **1939** | 1939_KS5-S244-A_for | ttcgcggcctcggccgacggcaccggtttcgcggagggcgcgggactgctcctcctggagcgg |
| **1940** | 1940_KS5-S244-A_rev | aggaggagcagtcccgcgccctccgcgaaaccggtgccgtcggccgaggccgcgaaagccttcg |
| **1941** | 1941_KS5-R221-E_Q222-L_for | acgcccggcgcgttcaccgagttctccgagctgggcgcgctgtctccggacggccgctcg |
| **1942** | 1942_KS5-R221-E_Q222-L_rev | agcggccgtccggagacagcgcgcccagctcggagaactcggtgaacgcgccgggcgtcg |
| **1943** | 1943_KS5-F243-M_S244-A_for | ttcgcggcctcggccgacggcaccggtatggcggagggcgcgggactgctcctcctggagcgg |
| **1944** | 1944_KS5-F243-M_S244-A_rev | aggaggagcagtcccgcgccctccgccataccggtgccgtcggccgaggccgcgaaagccttcg |
| **1945** | 1945_KS5-G214-T_for | tgtcaccgtgatgtcgacgcccaccgcgttcaccgagttctcccgccagggcgcgctg |
| **1946** | 1946_KS5-G214-T_rev | tggcgggagaactcggtgaacgcggtgggcgtcgacatcacggtgacaccgccgg |
| **1947** | 1947_KS5-G214-S_for | tgtcaccgtgatgtcgacgccctccgcgttcaccgagttctcccgccagggcgcgctg |
| **1948** | 1948_KS5-G214-S_rev | tggcgggagaactcggtgaacgcggagggcgtcgacatcacggtgacaccgccgg |
| **1949** | 1949_KS5-G214-A_for | tgtcaccgtgatgtcgacgcccgccgcgttcaccgagttctcccgccagggcgcgctg |
| **1950** | 1950_KS5-G214-A_rev | tggcgggagaactcggtgaacgcggcgggcgtcgacatcacggtgacaccgccgg |
| **1951** | 1951_KS5-A215-S_for | tgtcaccgtgatgtcgacgcccggctcgttcaccgagttctcccgccagggcgcgctg |
| **1952** | 1952_KS5-A215-S_rev | tggcgggagaactcggtgaacgagccgggcgtcgacatcacggtgacaccgccggccagcg |
| **1953** | 1953_KS5-A215-G_for | tgtcaccgtgatgtcgacgcccggcgggttcaccgagttctcccgccagggcgcgctg |
| **1954** | 1954_KS5-A215-G_rev | tggcgggagaactcggtgaacccgccgggcgtcgacatcacggtgacaccgccggccagcg |
| **1955** | 1955_KS5-A215-P_for | tgtcaccgtgatgtcgacgcccggcccgttcaccgagttctcccgccagggcgcgctg |
| **1956** | 1956_KS5-A215-P_rev | tggcgggagaactcggtgaacgggccgggcgtcgacatcacggtgacaccgccggccagcg |
| **1957** | 1957_KS5-G214-T_A215-S_for | tgtcaccgtgatgtcgacgcccacctcgttcaccgagttctcccgccagggcgcgctg |

| | | |
|---|---|---|
| **1958** | 1958_KS5-G214-T_A215-S_rev | tggcgggagaactcggtgaacgaggtgggcgtcgacatcacggtgacaccgccgg |
| **1959** | 1959_KS5-G214-S_A215-G_for | tgtcaccgtgatgtcgacgccctccgggttcaccgagttctcccgccagggcgcgctg |
| **1960** | 1960_KS5-G214-S_A215-G_rev | tggcgggagaactcggtgaacccggagggcgtcgacatcacggtgacaccgccgg |
| **1961** | 1961_KS5-G214-A_A215-P_for | tgtcaccgtgatgtcgacgcccgccccgttcaccgagttctcccgccagggcgcgctg |
| **1962** | 1962_KS5-G214-A_A215-P_rev | tggcgggagaactcggtgaacggggcgggcgtcgacatcacggtgacaccgccgg |
| **1965** | 1965_KS5_TGFS243-TGWA_for | ttcgcggcctcggccgacggcaccggttgggcggagggcgcgggactgctcctcctggagcgg |
| **1966** | 1966_KS5_TGFS243-TGWA_rev | tccaggaggagcagtcccgcgccctccgcccaaccggtgccgtcggccgaggccgcg |
| **1967** | 1967_KS5_TGFS243-TGWG_for | ttcgcggcctcggccgacggcaccggttgggggagggcgcgggactgctcctcctggagcgg |
| **1968** | 1968_KS5_TGFS243-TGWG_rev | tccaggaggagcagtcccgcgccctccccccaaccggtgccgtcggccgaggccgcg |
| **1969** | 1969_KS5_TGFS241-MVWG_for | aaggctttcgcggcctcggccgacggcatggtgtgggggagggcgcgggactgctcctcctggagcgg |
| **1970** | 1970_KS5_TGFS241-MVWG_rev | tccaggaggagcagtcccgcgccctccccccacaccatgccgtcggccgaggccgcgaaagccttcgagcgg |
| **1971** | 1971_KS5_TGFS241-FGPS_for | aaggctttcgcggcctcggccgacggcttcggtccctcggagggcgcgggactgctcctcctggagcgg |
| **1972** | 1972_KS5_TGFS241-FGPS_rev | aggaggagcagtcccgcgccctccgagggaccgaagccgtcggccgaggccgcgaaagccttcgagcgg |

#### Fermentation, extraction and analysis

##### Fermentation for LC-MS

Agar plugs (1 cm²) with colonies were inoculated into 24-well plates containing 2 mL TSB and five glass beads and incubated at 30 °C, 180 rpm for 2 days as precultures. Aliquots (750 µL) were used to inoculate 15 mL SM16 medium supplemented with 20 g/L XAD16 resin and one glass bead in 250 mL Erlenmeyer flasks (5% v/v inoculum). Cultures were grown for 5 days at 30 °C, 180 rpm with foam stoppers and a 5 cm throw. Remaining precultures were stored at –80 °C as glycerol stocks.

##### Extraction of premonensin and derivatives

Resin and cell paste were harvested by centrifugation (25 min, 3900 rpm, 4 °C), stored at –80 °C overnight, then extracted with 6 mL ethyl acetate and glass beads after vortexing (30 s) and incubation at 19 °C, 180 rpm overnight. Samples were centrifuged (10 min, 3900 rpm, 4 °C), and the organic phase was evaporated at 38 °C. Residues were reconstituted in 3 mL LC–MS grade acetonitrile, transferred to microcentrifuge tubes, frozen (–20 °C), and centrifuged (30 min, 3900 rpm, 4 °C). Supernatants were transferred to HPLC vials, and 2 µL (A495 variants) or 5 µL ($DH4^0$ variant) were injected for HPLC–MS analysis.

##### Large-scale fermentation for qNMR

Agar plugs (3 cm²) were added to 15 mL TSB and the culture was grown for 2 days at 30 °C, 180 rpm. Precultures (75 mL) were used to inoculate 1.5 L SM16 medium supplemented with 20 g/L XAD16 resin. Cultivation was carried out for 5 days under the same conditions, followed by centrifugation (25 min, 3900 rpm, 4 °C) and storage at –80 °C.

##### Premonensin extraction for qNMR

Resin–cell paste was snap-frozen in liquid nitrogen, ground, and freeze-dried. Extraction was performed overnight at 19 °C, 180 rpm with glass beads (1.7–2.1 mm) and ethyl acetate (1 g sample : 5 g beads : 20 mL solvent), followed by two 1 h extractions with 300 mL ethyl acetate each. Supernatants were collected by centrifugation (20 min, 3900 rpm, 4 °C), pooled, dried over $MgSO_4$, and concentrated under reduced pressure at 30 °C. Crude extracts were dissolved in minimal ethyl acetate, and fractionated on a silica column with sequential elution using cyclohexane/ethyl acetate (500 mL 80:20, 500 mL 60:40, 500 mL 50:50). Fractions containing premonensin were verified by TLC with vanillin staining, then pooled and concentrated under vacuum.

#### HPLC-MS Analysis

Fermentation extracts were analyzed by HPLC–MS for screening and verified by repeated HPLC–MS/MS measurements. Data processing was performed in MZmine3 as previously described [3].

### HPLC method

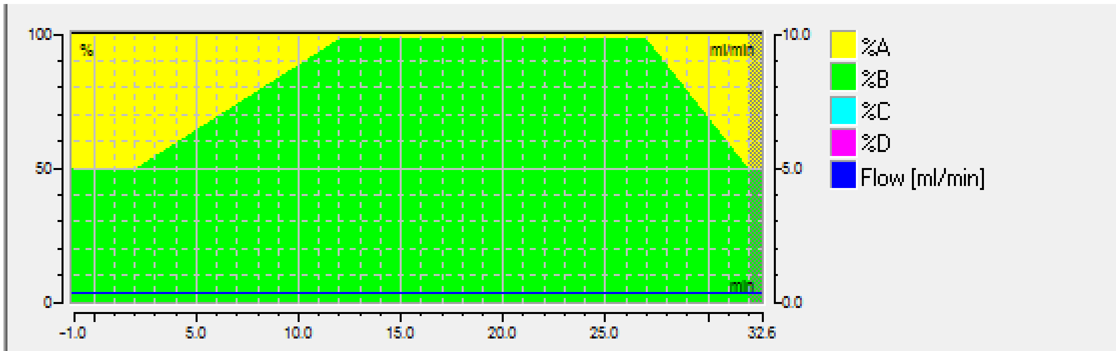


| | Retention [min] | Flow [ml/min] | %B | %C | %D | Curve |
|---|---|---|---|---|---|---|
| 1 | -1.000 | 0.300 | 50.0 | 0.0 | 0.0 | |
| 2 | 2.000 | 0.300 | 50.0 | 0.0 | 0.0 | 5 |
| 3 | 12.000 | 0.300 | 98.0 | 0.0 | 0.0 | 5 |
| 4 | 27.000 | 0.300 | 98.0 | 0.0 | 0.0 | 5 |
| 5 | 32.000 | 0.300 | 50.0 | 0.0 | 0.0 | 5 |

Insert Line
Delete Line
Clean Up

Figure S 5: HPLC method used in this study. The mobile phase consisted of water with 0.1% formic acid (A) and acetonitrile with 0.1% formic acid (B). Internal calibration was performed using a lock mass of 622.028960 m/z (Hexakis(1H,1H,2H-perfluoroethoxy)phosphazene) and sodium formate clusters.

### MS Method

#### Method Set: D:\Methods\Susanna\110-1300 autoMSMS pos_.m

#### otofControl

##### General Information

**Method Name:** 110-1300 autoMSMS pos_.m
**Application Name:** Bruker otofControl
**Device Type:** compact
**Operator:** Demo User
**Operating System:** Windows 7 Professional

**Saved:** 2018/09/05 09:38:08+02:00
**Application Version:** 4.1.3.5
**Device Serial Number:** 8255754.20147
**Host:** COMPACT-20147
**Organisation:** Bruker Daltonik GmbH

##### Chromatogram

###### Chromatogram Traces

| Enabled | Color | Type | Masses | Width | Polarity | Filter |
|---|---|---|---|---|---|---|
| On | Red | BPC | | | ± | MS |
| On | Blue | TIC | | | ± | MS |
| On | Black | TIC | | | ± | All MS/MS |

##### SPL

**Scheduled Precursor List:** Off

##### Segment 1

##### 0 .... 0.02 min

###### Main

**Polarity:** Positive
**Mass Range from:** 110 m/z
**Rolling Average:** Off
**Spectra rate:** 8.00 Hz
**Scan Mode:** MS
**Mass Range to:** 1300 m/z
**Rolling Average No.:** 2
**View:** Expert

###### Mode

**Save Spectra:** Line and Profile Spectra
**Absolute Threshold (per 1000 sum.):** 25 cts.
**Mark as Calibration Segment:** Off
**Line Spectra Calculation:** Use Maximum Intensity
**Peak Summation Width:** 3 pts.
**Focus Active:** Off

###### Source

**Source:** ESI
**End Plate Offset:** 500 V
**Nebulizer:** 2.2 Bar
**Dry Temp:** 220 °C
**Capillary:** 4500 V
**Dry Gas:** 10.0 l/min
**Divert Valve:** Waste 1-6

##### Tune

| | | | |
|---|---|---|---|
| **Funnel 1 RF:** | 150.0 Vpp | **Funnel 2 RF:** | 200.0 Vpp |
| **isCID Energy:** | 0.0 eV | **Hexapole RF:** | 50.0 Vpp |
| **Ion Energy:** | 4.0 eV | **Low Mass:** | 90.0 m/z |
| **Collision Energy:** | 7.0 eV | **Pre Pulse Storage:** | 5.0 µs |
| **Stepping:** | On | **Mode:** | Basic |
| **Collision RF from:** | 550.0 Vpp | **Collision RF to:** | 550.0 Vpp |
| **Transfer Time from:** | 80.0 µs | **Transfer Time to:** | 80.0 µs |
| **Timing from:** | 50 % | **Timing to:** | 50 % |
| **Collision Energy from:** | 100 % | **Collision Energy to:** | 250 % |
| **Timing from:** | 50 % | **Timing to:** | 50 % |

### MS/MS

| | |
|---|---|
| **Auto MS/MS:** | Off |

##### *MRM*

| | |
|---|---|
| **MRM:** | Off |

##### *isCID*

| | |
|---|---|
| **isCID (MS-MS/MS):** | Off |

##### *bbCID*

| | |
|---|---|
| **bbCID (MS-MS/MS):** | Off |

#### Segment 2
#### 0.02 .... 0.3 min

##### Main

| | | | |
|---|---|---|---|
| **Polarity:** | Positive | **Scan Mode:** | MS |
| **Mass Range from:** | 110 m/z | **Mass Range to:** | 1300 m/z |
| **Rolling Average:** | Off | **Rolling Average No.:** | 2 |
| **Spectra rate:** | 8.00 Hz | **View:** | Expert |

##### Mode

| | | | |
|---|---|---|---|
| **Save Spectra:** | Line and Profile Spectra | **Line Spectra Calculation:** | Use Maximum Intensity |
| **Absolute Threshold (per 1000 sum.):** | 25 cts. | **Peak Summation Width:** | 3 pts. |
| | | **Focus Active:** | Off |
| **Mark as Calibration Segment:** | On | | |

##### Source

| | | | |
|---|---|---|---|
| **Source:** | ESI | | |
| **End Plate Offset:** | 500 V | **Capillary:** | 4500 V |
| **Nebulizer:** | 2.2 Bar | **Dry Gas:** | 10.0 l/min |
| **Dry Temp:** | 220 °C | **Divert Valve:** | Source 1-2 |

##### Tune

| | | | |
|---|---|---|---|
| **Funnel 1 RF:** | 150.0 Vpp | **Funnel 2 RF:** | 200.0 Vpp |
| **isCID Energy:** | 0.0 eV | **Hexapole RF:** | 50.0 Vpp |
| **Ion Energy:** | 4.0 eV | **Low Mass:** | 90.0 m/z |
| **Collision Energy:** | 7.0 eV | **Pre Pulse Storage:** | 5.0 µs |
| **Stepping:** | On | **Mode:** | Basic |
| **Collision RF from:** | 550.0 Vpp | **Collision RF to:** | 550.0 Vpp |
| **Transfer Time from:** | 80.0 µs | **Transfer Time to:** | 80.0 µs |
| **Timing from:** | 50 % | **Timing to:** | 50 % |
| **Collision Energy from:** | 100 % | **Collision Energy to:** | 250 % |
| **Timing from:** | 50 % | **Timing to:** | 50 % |

### MS/MS

| | |
|---|---|
| **Auto MS/MS:** | Off |

##### *MRM*

| | |
|---|---|
| **MRM:** | Off |

##### *isCID*

| | |
|---|---|
| **isCID (MS-MS/MS):** | Off |

##### *bbCID*

| | |
|---|---|
| **bbCID (MS-MS/MS):** | Off |

#### Segment 3
#### 0.3 .... unlimited min

##### Main

| | | | |
|---|---|---|---|
| **Polarity:** | Positive | **Scan Mode:** | Auto MS/MS |
| **Mass Range from:** | 110 m/z | **Mass Range to:** | 1300 m/z |
| **Rolling Average:** | Off | **Rolling Average No.:** | 2 |
| **Spectra rate:** | 8.00 Hz | **View:** | Expert |

##### Mode

| | | | |
|---|---|---|---|
| **Save Spectra:** | Line and Profile Spectra | **Line Spectra Calculation:** | Use Maximum Intensity |
| **Absolute Threshold (per 1000 sum.):** | 25 cts. | **Peak Summation Width:** | 3 pts. |
| | | **Focus Active:** | Off |
| **Mark as Calibration Segment:** | Off | | |

##### Source

| | | | |
|---|---|---|---|
| **Source:** | ESI | | |
| **End Plate Offset:** | 500 V | **Capillary:** | 4500 V |
| **Nebulizer:** | 2.2 Bar | **Dry Gas:** | 10.0 l/min |
| **Dry Temp:** | 220 °C | **Divert Valve:** | Waste 1-6 |

#### Tune

| | | | |
|---|---|---|---|
| **Funnel 1 RF:** | 150.0 Vpp | **Funnel 2 RF:** | 200.0 Vpp |
| **isCID Energy:** | 0.0 eV | **Hexapole RF:** | 50.0 Vpp |
| **Ion Energy:** | 4.0 eV | **Low Mass:** | 90.0 m/z |
| **Collision Energy:** | 7.0 eV | **Pre Pulse Storage:** | 5.0 µs |
| **Stepping:** | On | **Mode:** | Basic |
| **Collision RF from:** | 550.0 Vpp | **Collision RF to:** | 550.0 Vpp |
| **Transfer Time from:** | 80.0 µs | **Transfer Time to:** | 80.0 µs |
| **Timing from:** | 50 % | **Timing to:** | 50 % |
| **Collision Energy from:** | 100 % | **Collision Energy to:** | 250 % |
| **Timing from:** | 50 % | **Timing to:** | 50 % |

## MS/MS

| | | | |
|---|---|---|---|
| **Auto MS/MS:** | On | | |
| **Precursor Ion List:** | Exclude | **Cycle Time:** | 0.5 sec |
| **Threshold (per 1000 sum.) Absolute:** | 400 cts | **Active Exclusion:** | On |
| | | **Exclude after:** | 3 Spectra |
| | | **Reconsider Precursor:** | On |
| **Release after:** | 0.20 min. | **Smart Exclusion:** | Off |
| **if Curent Intens./Prev. Intens.:** | 1.8 | | |
| **Smart Exclusion:** | 2 x | | |

##### Exclude Mass List

| Mass Range Start | Mass Range End | |
|---|---|---|
| 102.08 | 102.18 | 1 |
| 621.98 | 622.08 | 2 |
| 643.96 | 644.06 | 3 |
| 659.94 | 660.04 | 4 |

##### Auto MS/MS Preference

| | | | |
|---|---|---|---|
| **Preferred Range:** | Off | | |
| **Preferred Range Low:** | 2 | **Preferred Range High:** | 5 |
| **Exclude Singly:** | Off | **Exclude unknown:** | Off |
| **Group Length:** | 3 | **Strict Active Exclusion:** | Off |
| **Sort Precursors by:** | Intensity | **Preferred mass list:** | Empty list |

##### Auto MS/MS Multi CE

**Auto MS/MS Multi CE:** Off

##### SILE

**SILE:** Off

##### CID

**Fallback Charge State:** 1 z

###### Isolation + Fragmentation List

| Type | Mass [m/z] | Width [m/z] | Collision Energy [eV] | Charge State | |
|---|---|---|---|---|---|
| Base | 100.00 | 4.00 | 20.0 | 1 | 1 |
| Base | 500.00 | 5.00 | 20.0 | 1 | 2 |
| Base | 1000.00 | 6.00 | 20.0 | 1 | 3 |
| Base | 1300.00 | 8.00 | 30.0 | 1 | 4 |

##### CID Acquisition

| | | | |
|---|---|---|---|
| **Acquisition:** | On | **Spectra Rate MS:** | 8.00 Hz |
| **MS/MS low (per 1000 sum.):** | 10000.0 cts. | **MS/MS low:** | 1600 x |
| **MS/MS high:** | 100000.0 cts. | **MS/MS high:** | 800 x |
| **Total Cycle Time Range:** | n/a sec | **Absolute Threshold :** | n/a cts. |

##### MRM

**MRM:** Off

##### isCID

**isCID (MS-MS/MS):** Off

##### bbCID

**bbCID (MS-MS/MS):** Off

#### MS$^1$ and MS$^2$ data of PreA/B, KR4$^0$PreA/B, and DH4$^0$PreA/B

Table S 2 List of characteristic adducts of PreA/B, KR4$^0$PreA/B, and DH4$^0$PreA/B

| | **meas. *m/z*** | **ion formula** | **calc. *m/z*** | **err [ppm]** |
|---|---|---|---|---|
| **PreB** | 561.4168 | $C_{34}H_{57}O_6$ | 561.4150 | 3.2 |
| | 578.4422 | $C34H_{60}NO_6$ | 578.4415 | 1.2 |
| | 583.3978 | $C_{34}H_{56}NaO_6$ | 583.3969 | 1.5 |
| | 599.3730 | $C_{34}H_{56}KO_6$ | 599.3708 | 3.7 |
| **PreA** | 575.4303 | $C_{35}H_{59}O6$ | 575.4306 | -0.5 |
| | 592.4568 | $C_{35}H_{62}NO6$ | 592.4572 | -0.7 |
| | 597.4139 | $C_{35}H_{58}NaO_6$ | 597.4126 | 2.2 |
| | 613.3832 | $C_{35}H_{58}KO_6$ | 613.3865 | -5.4 |
| **KR4$^0$PreB** | 575.3941 | $C_{34}H_{55}O_7$ | 575.3942 | -0.2 |
| | 592.4215 | $C_{34}H_{58}NO_7$ | 592.4208 | 1.2 |
| | 597.3765 | $C_{34}H_{54}NaO_7$ | 597.3762 | 0.5 |
| | 613.3491 | $C_{34}H_{54}KO_7$ | 613.3501 | -1.6 |
| **KR4$^0$PreA** | 589.4113 | $C_{35}H_{57}O_7$ | 589.4099 | 2.4 |
| | 606.4353 | $C_{35}H_{50}NO_7$ | 606.4364 | -1.8 |
| | 611.3926 | $C_{35}H_{56}NaO_7$ | 611.3918 | 1.3 |
| | 627.3668 | $C_{35}H_{56}KO_7$ | 627.3662 | 1.0 |
| **DH4$^0$PreB** | 594.4351 | $C_{34}H_{60}NO_7$ | 594.4365 | -2.4 |
| | 599.3915 | $C_{34}H_{56}NaO_7$ | 599.3918 | -0.5 |
| | 615.3701 | $C_{34}H_{56}KO_7$ | 615.3659 | 6.8 |
| **DH4$^0$PreA** | 608.4575 | $C_{35}H_{62}NO_7$ | 608.4521 | 8.8 |
| | 613.4079 | $C_{35}H_{58}NaO_7$ | 613.4075 | 0.7 |

**PreA/B**

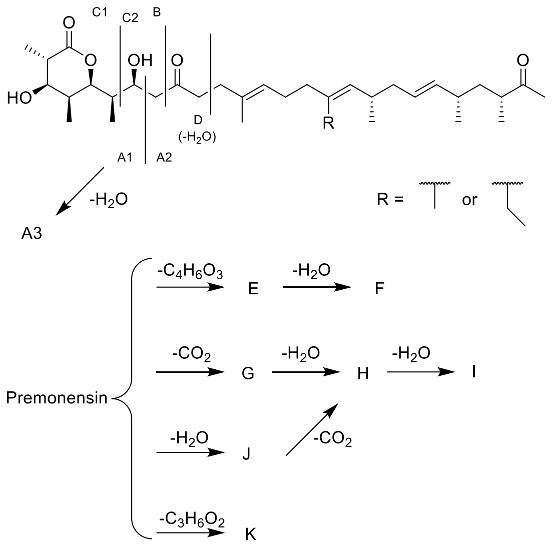


Figure S 6 Premonensin A/B structures and fragmentation patterns.

Table S 3 Mass spectrometric data of the fragment ions of PreA/B.

| Fragment | meas. *m/z* | ion formula | *m/z* | err [ppm] |
|---|---|---|---|---|
| | | **PreB** | | |
| **C1** | 193.0854 | $C_9H_{14}NaO_3$ | 193.0835 | 9.8 |
| **A3** | 205.0846 | $C_{10}H_{14}NaO_3$ | 205.0835 | 5.4 |
| **A1** | 223.095 | $C_{10}H_{16}NaO_4$ | 223.0941 | 4.0 |
| **B** | 237.1092 | $C_{11}H_{18}NaO_4$ | 237.1097 | -2.1 |
| **D** | 263.1271 | $C_{13}H_{20}NaO_4$ | 263.1254 | 6.5 |
| **A2** | 383.2947 | $C_{24}H_{40}NaO_2$ | 383.2921 | 6.8 |
| **C2** | 411.2864 | $C_{25}H_{40}NaO_3$ | 411.2870 | -1.5 |
| **F** | 463.3529 | $C_{30}H_{48}NaO_2$ | 463.3547 | -3.9 |
| **E** | 481.3675 | $C_{30}H_{50}NaO_3$ | 481.3652 | 4.8 |
| **I** | 503.3856 | $C_{33}H_{52}NaO_2$ | 503.3860 | -0.8 |
| **K** | 509.3608 | $C_{31}H_{50}NaO_4$ | 509.3601 | 1.4 |
| **J** | 521.3985 | $C_{33}H_{54}NaO_3$ | 521.3965 | 3.8 |
| **G** | 539.4011 | $C_{33}H_{56}NaO_4$ | 539.4071 | -11.1 |
| **H** | 565.3904 | $C_{34}H_{54}NaO_5$ | 565.3863 | 7.3 |
| | | **PreA** | | |
| **C1** | 193.0839 | $C_9H_{14}NaO_3$ | 193.0835 | 2.2 |
| **A3** | 205.0837 | $C_{10}H_{14}NaO_3$ | 205.0835 | 1.0 |
| **A1** | 223.0944 | $C_{10}H_{16}NaO_4$ | 223.0941 | 1.3 |
| **B** | 237.1112 | $C_{11}H_{18}NaO_4$ | 237.1097 | 6.2 |
| **C2** | 263.1261 | $C_{13}H_{20}NaO_4$ | 263.1254 | 2.8 |
| **A2** | 397.3071 | $C_{25}H_{42}NaO_2$ | 397.3077 | -1.4 |

| | | | | |
|---|---|---|---|---|
| **D** | 425.3012 | $C_{26}H_{42}NaO_3$ | 425.3026 | -3.2 |
| **F** | 477.3704 | $C_{31}H_{50}NaO_2$ | 477.3703 | 0.2 |
| **E** | 495.3827 | $C_{31}H_{52}NaO_3$ | 495.3809 | 3.6 |
| **I** | 517.4014 | $C_{34}H_{54}NaO_2$ | 517.4016 | -0.4 |
| **K** | 523.3772 | $C_{32}H_{52}NaO_4$ | 523.3758 | 2.7 |
| **J** | 535.4060 | $C_{34}H_{56}NaO_3$ | 535.4122 | -11.5 |
| **G** | 553.4148 | $C_{34}H_{58}NaO_4$ | 553.4227 | -14.3 |
| **H** | 579.3977 | $C_{35}H_{56}NaO_5$ | 579.4020 | -7.5 |

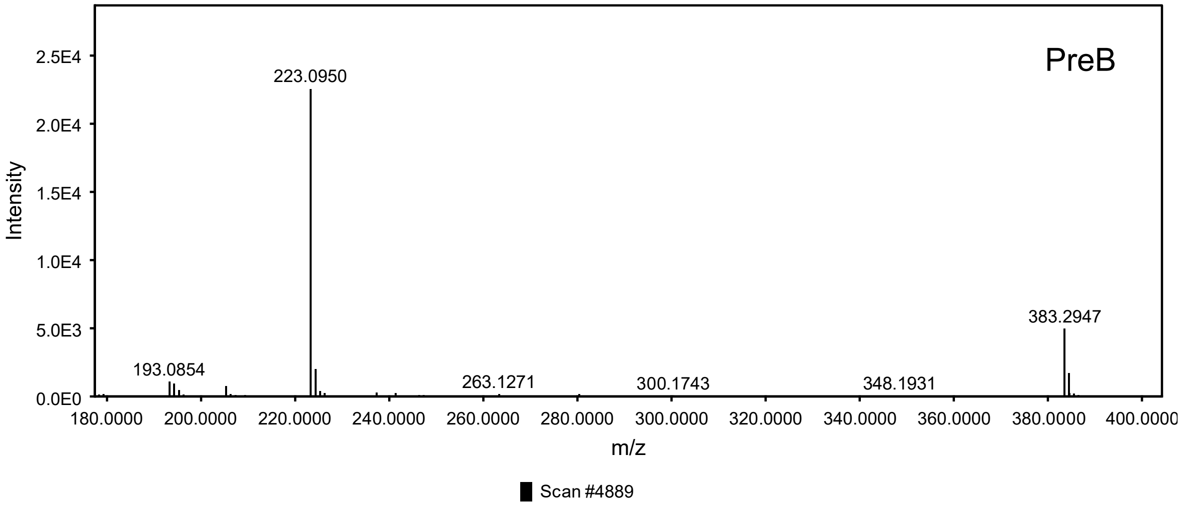


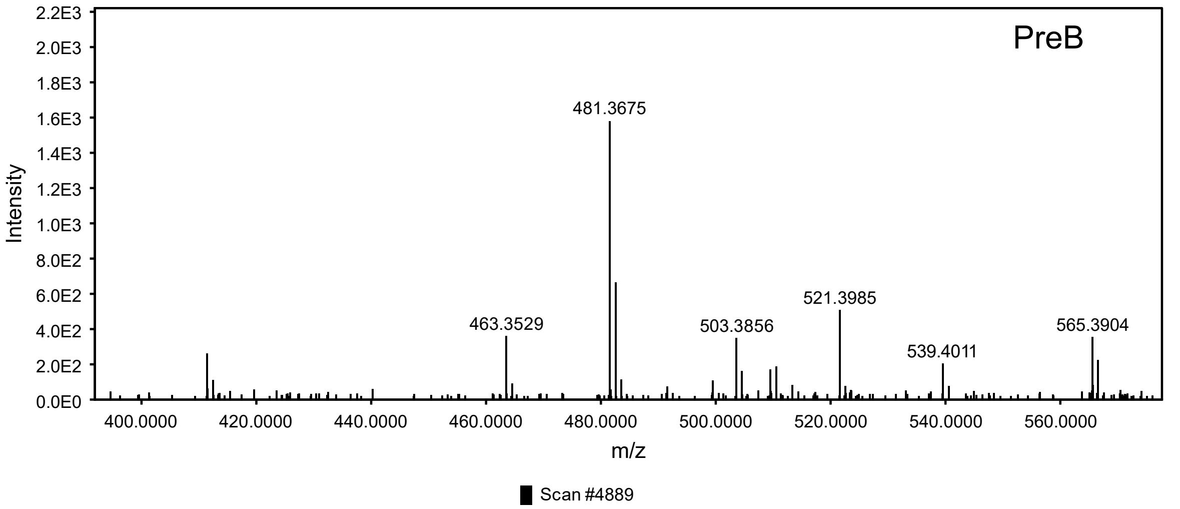

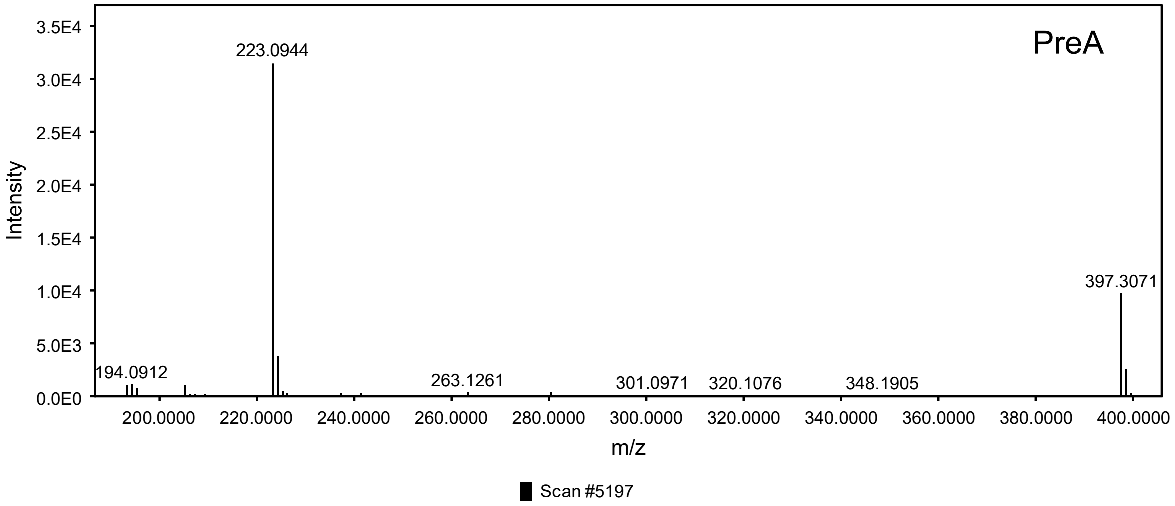


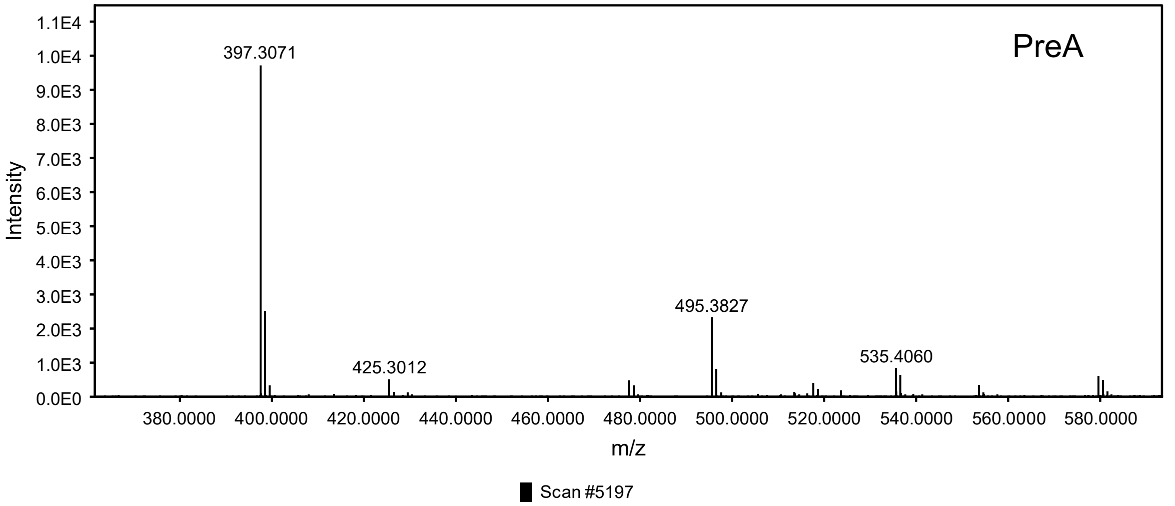


Figure S 7 $MS^2$ spectra of PreA/B.

**KR4$^0$PreA/B**

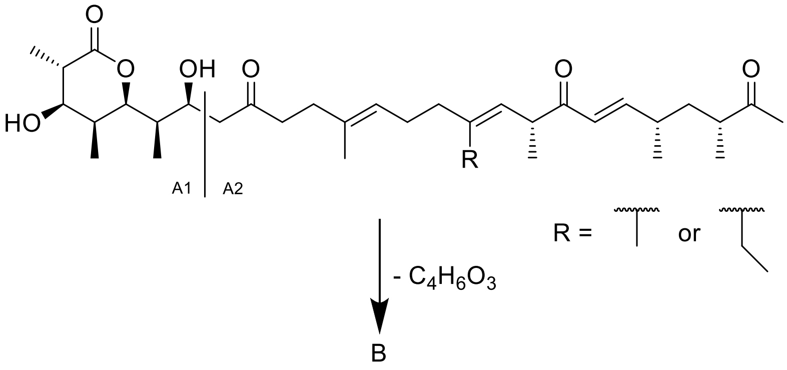


Figure S 8 KR4$^0$PreA/B structures and fragmentation patterns.

Table S 4 Mass spectrometric data of the fragment ions of KR4$^0$PreA/B.

| Fragment | meas. m/z | ion formula | m/z | err [ppm] |
|---|---|---|---|---|
| | | **KR4$^0$PreA/B** | | |
| **A1** | 223.0952 | $C_{10}H_{16}NaO_4$ | 223.0941 | 4.9 |
| **A2** | 397.2723 | $C_{24}H_{38}NaO_3$ | 397.2713 | 2.5 |
| **B** | 495.3452 | $C_{30}H_{48}NaO_4$ | 495.3445 | 1.4 |
| | | **KR4$^0$PreA/B** | | |
| **A1** | 223.0932 | $C_{10}H_{16}NaO_4$ | 223.0941 | -4.0 |
| **A2** | 411.2853 | $C_{25}H_{40}NaO_3$ | 411.2870 | -4.1 |
| **B** | 509.3599 | $C_{31}H_{50}NaO_4$ | 509.3601 | -0.4 |

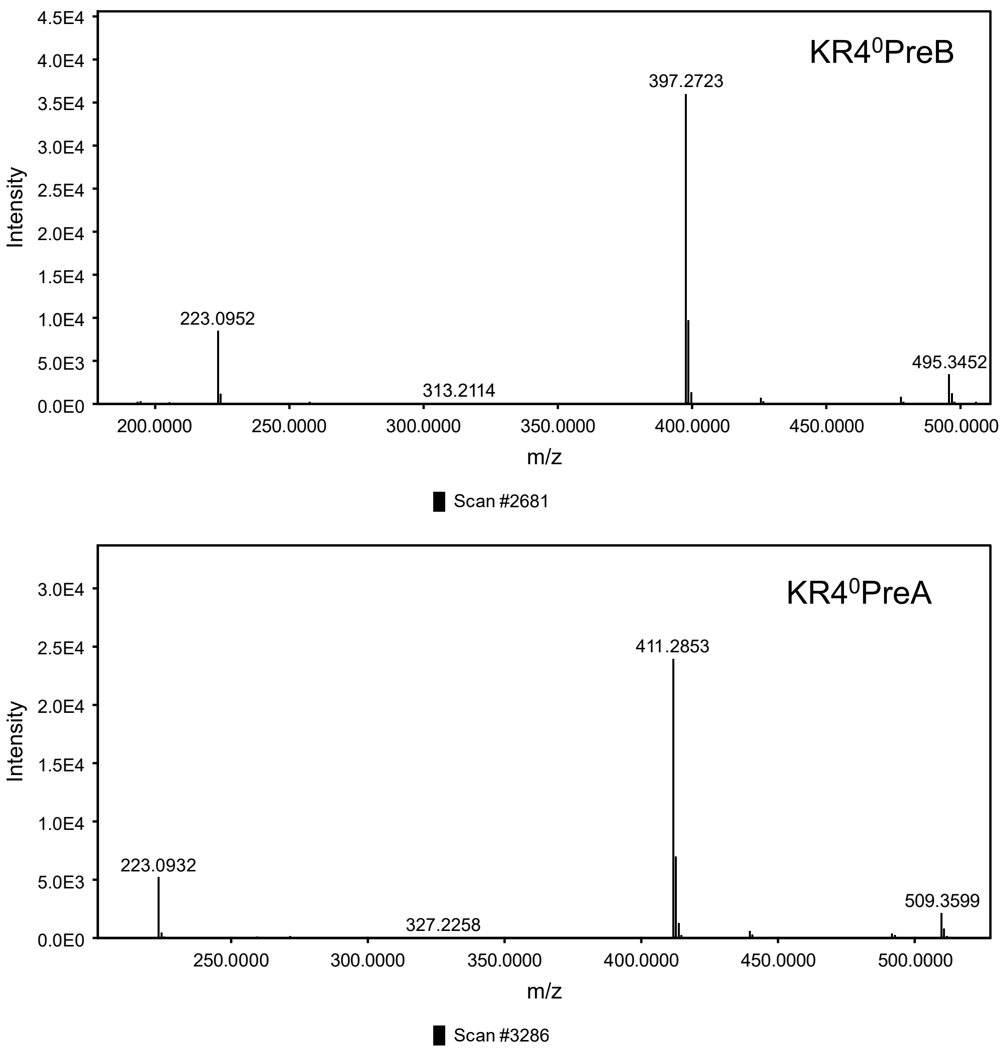


Figure S 9 $MS^2$ spectra of $KR4^0$PreA/B.

**DH4⁰PreA/B**

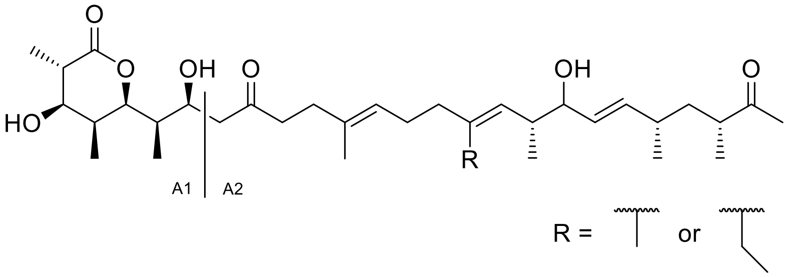


Figure S 10 DH4⁰PreA/B structures and fragmentation patterns.

Table S 5 Mass spectrometric data of the fragment ions of DH4⁰PreA/B.

| Fragment | meas. m/z | ion formula | m/z | err [ppm] |
|---|---|---|---|---|
| | | **DH4⁰PreA/B** | | |
| **A1** | 223.0938 | $C_{10}H_{16}NaO_4$ | 223.0941 | -1.3 |
| **A2** | 399.2857 | $C_{24}H_{40}NaO_3$ | 399.2870 | -3.3 |
| | | **DH4⁰PreA/B** | | |
| **A1** | 223.0942 | $C_{10}H_{16}NaO_4$ | 223.0941 | 0.4 |
| **A2** | 413.3023 | $C_{25}H_{42}NaO_3$ | 413.3026 | -0.7 |

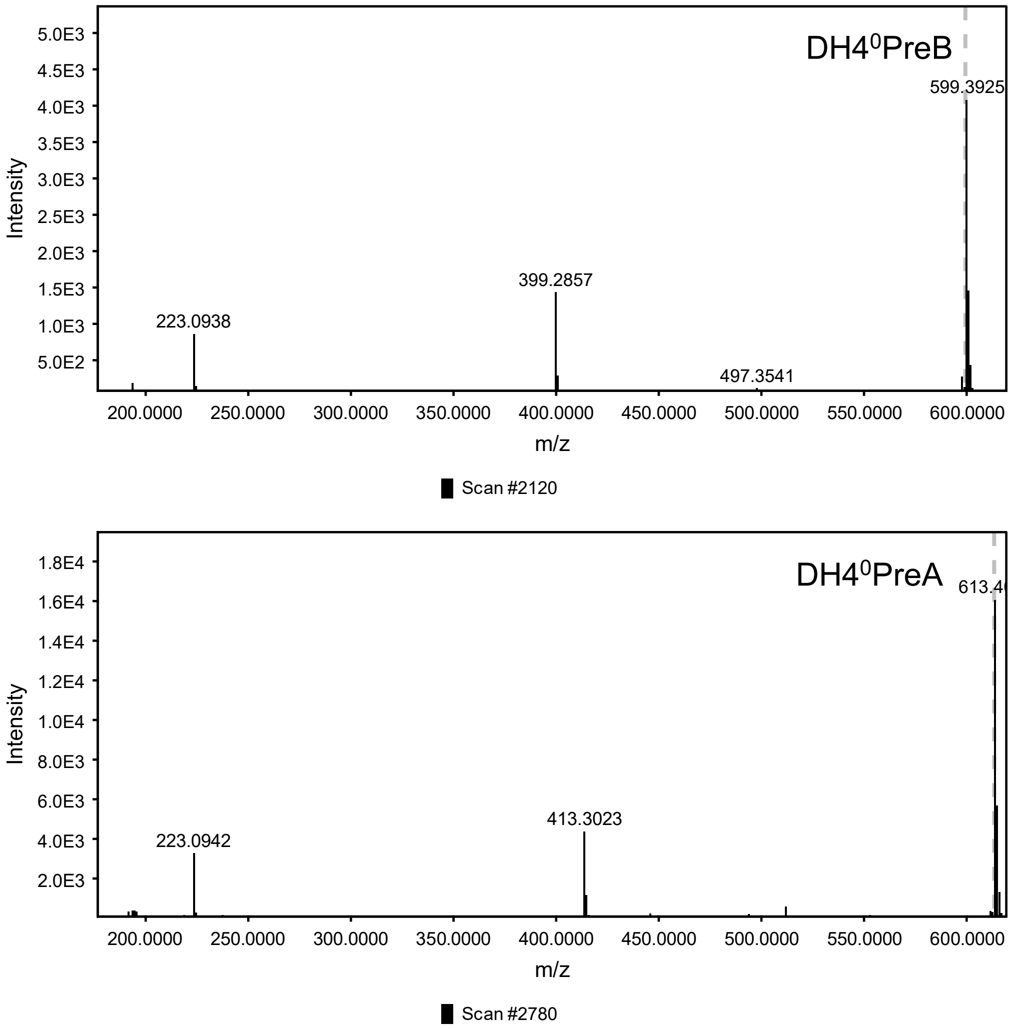


Figure S 11 $MS^2$ spectra of $DH4^0$PreA/B.
